# Hippocampal cells multiplex positive and negative engrams

**DOI:** 10.1101/2020.12.11.419887

**Authors:** Monika Shpokayte, Olivia McKissick, Xiaonan Guan, Bingbing Yuan, Bahar Rahsepar, Fernando R. Fernandez, Evan Ruesch, Stephanie L. Grella, John A. White, X. Shawn Liu, Steve Ramirez

## Abstract

The hippocampus is involved in processing a variety of mnemonic computations specifically the spatiotemporal components and emotional dimensions of contextual memory.^1–3^ Recent studies have demonstrated vast structural and functional heterogeneity along the dorsal-ventral axis^1, 5^ of the hippocampus. The ventral hippocampus has been shown to be important in the processing of emotion and valence.^6–9^ Here, we combine transgenic and all-virus based activity-dependent tagging strategies to visualize multiple valence-specific engrams in the vHPC and demonstrate two partially segregated cell populations and projections that respond to appetitive and aversive experiences. Next, using RNA sequencing and DNA methylation sequencing approaches, we find that vHPC appetitive and aversive engram cells display distinct transcriptional programs and DNA methylation landscapes compared to a neutral engram population. Additionally, while optogenetic manipulation of tagged cell bodies in vHPC is not sufficient to drive appetitive or aversive behavior in real-time place preference, stimulation of tagged vHPC terminals projecting to the amygdala and nucleus accumbens (NAc), but not the prefrontal cortex (PFC), had the capacity drive preference and avoidance. These terminals can also undergo a “switch” or “reset” in their capacity to drive either, thereby demonstrating their adaptable contributions to behavior. We conclude that the vHPC contains genetically, cellularly, and behaviorally distinct populations of cells processing appetitive and aversive memory engrams. Together, our findings provide a novel means by which to visualize multiple engrams within the same brain and point to their unique genetic signatures as reference maps for the future development of new therapeutic strategies.

**One sentence summary:** The hippocampus contains neurons that correspond to positive and negative engrams, which are segregated by their molecular, cellular, and projection-specific features.

---

Within the brain we find a rich repository of memories that can be imbued with positive and negative valenced information. These experiences leave enduring structural and functional changes that are parceled up into discrete sets of cells and circuits comprising the memory engram^10^. Recent studies have successfully visualized and manipulated defined sets of cells previously active during a single experience^10–17^. However, how multiple engrams of varying valences (hereafter defined as cells that differentially respond to appetitive or aversive events in a stimulus-independent manner^18^) are represented within the same brain region remains poorly understood. Previous studies have suggested the ventral hippocampus (vHPC) selectively demarcates and relays emotional information to various downstream targets. We sought to characterize the molecular and cellular identities and the behaviorally-relevant functions of vHPC cells processing appetitive and aversive engrams, with a focus on ventral CA1 (vCA1). In order to address the question of how the vCA1 processes multiple emotional experiences, we first devised a strategy combining two cFos-based tagging methods with endogenous cfos immunohistochemistry to visualize engrams across three discrete timepoints. First, we anatomically charted the projection patterns of appetitively and aversively-tagged vCA1 cells to measure structural overlap and segregation across various downstream targets. Second, we performed genome-wide RNA sequencing (RNA-seq) and DNA methylation sequencing to investigate the genetic features of these sets of cells. Lastly, we behaviorally tested the causal role and functional flexibility vHPC cells in both a cell body and projection-specific manner.

First, to access cells across multiple timepoints and in an activity-dependent, within-subject manner, we used the Fos-based transgenic animal, TRAP2^18^, under the control of 4-Hydroxytamoxifen (4-OHT) paired with an all-virus Fos-based strategy under the control of doxycycline^10^ (Dox) (**Fig. 1a**). TRAP2+ mice expressing iCre-ERT2 recombinase, when injected with DMSO instead of 4-OHT, show no TdTomato expression (**Extended Data Fig. 1**), were injected bilaterally with a cocktail of viruses: AAV9-Flex-DIO-TdTomato and AAV-cFos-tTA + AAV-TRE-EYFP. The cfos-tTA strategy couples the *c-Fos* promoter to the tetracycline transactivator (tTA), which, in its protein form, directly binds to the tetracycline response element (TRE) in a doxycycline-(Dox)-dependent manner and drives the expression of a protein of interest (i.e. EYFP). Combining these two independent systems yields two inducible windows for tagging vHPC cells within the same subject (See Methods).

**Figure 1.**
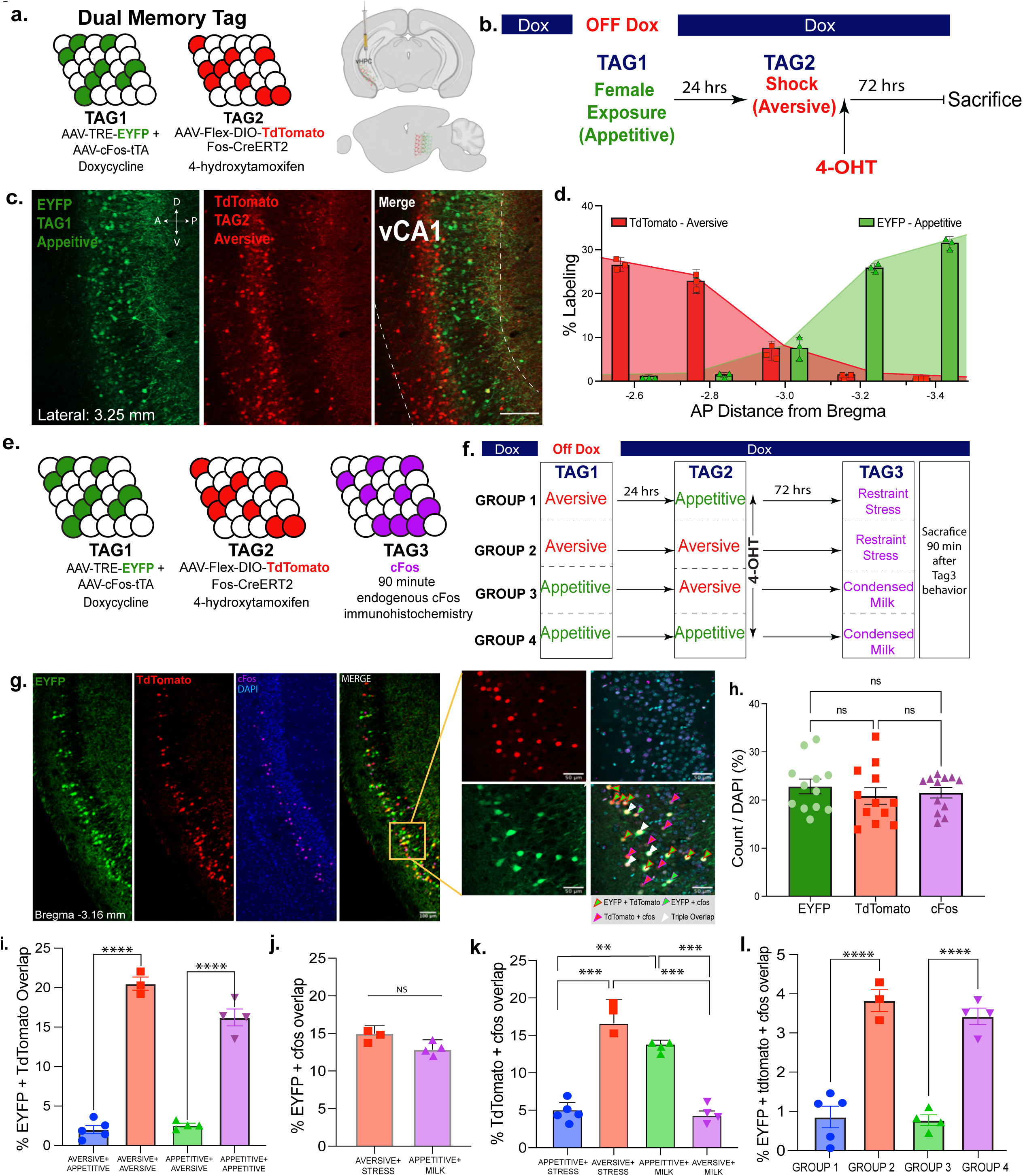
Hippocampus cells processing appetitive or aversive memory engrams are preferentially reactivated by their respective valences. **a**, Fos-CreERT2 mice were injected with a viral cocktail of 1:1 ratio of AAV9-c-Fos-tTA and AAV9-TRE-EYFP mixed in at 1:1 with AAV-Flex-DIO-TdTomato into the vHPC to create the ability to open two windows of tagging in the mouse. **b,** Schematic of the timeline for TAG1 and TAG2 which are doxycycline and 4-hydroxytamoxifen dependent, respectively. The aversive tag consists of 4 foot shocks and the appetitive tag consists of male to female exposure. **c,** Representative image of dual memory segregation in a sagittal section of the vHPC, specifically vCA1. **d,** Quantification (%Labeling) of EYFP+ (aversive) and TdTomato+ (appetitive) cells over DAPI along the anterior-posterior axis of vCA1 showing the segregation of valence. (N=3) **e,** Schematic to visualize cells active during three different behavioral experiences. TAG1 is doxycycline dependent in EYFP, TAG2 is 4-OHT dependent in TdTomato, and lastly, TAG3 is visualized with immunohistochemistry by staining for endogenous cfos 90 minutes after a behavioral experience. **f,** Schematic of groups for visualizing triple memory overlaps (n=4-6 per group). Groups were counterbalanced to avoid potential timeline biases **g,** Representative images of triple memory overlaps in vCA1 with a 20x zoom in showing the different representative overlaps. **h**, The number of tagged neurons in vCA1 are similar between EYFP, TdTomato, and cfos; there is no statistical difference between recruitment. (Multiple comparisons one-way Anova; N=10-12) **i,** Percent overlap for respective behavioral tags for EYFP + Tdtomato for aversive + appetitive (Group 1), aversive + aversive ( Group 2), appetitive + aversive ( Group 3), and appetitive + appetitive (Group4), (F (3, 12) = 170.0, P<0.0001). **j,** NS difference in cfos+EYFP overlapping cells in restraint stress + shock vs. sweetened condensed milk + female exposure. **k,** similar valences aversive and stress and appetitive and milk highly overlap with one another when compared to overlap of different valences, appetitive and stress and aversive and milk. Multiple comparisons one-way ANOVA; F (3, 12) = 20.77, P<0.0001. **l,** Triple overlap counts for EYFP + TdTomato + cfos (F (3, 12) = 10.55, P=0.0011).

After injecting the aforementioned virus cocktail into the vHPC, we used this “dual tagging” system to label cells processing appetitive or aversive engrams. Ten days after surgery and recovery, mice were taken off their DOX chow diet for 48 hours to open the first window of tagging. While off DOX, to tag the appetitive experience, the mice were subjected to female exposure for 1 hour in a clean homecage^13, 20^ and placed immediately back on the DOX diet for the remainder of the study. To tag the aversive experience, the second set of cells, 24 hours later male mice were subjected to four 2 second and 1.5 mA foot shocks. 30 minutes after the end of fear conditioning, the mice were injected with 40 mg/kg of 4-OHT and left undisturbed for 72 hours (**Fig. 1b**). We first observed a striking anatomical distinction between appetitive and aversive engrams along the anterior and posterior axis in sagittal sections of vCA1 (**Fig. 1c**). Appetitive engram cells were predominantly located in more posterior sections, whereas, aversive engram cells were largely present in anterior sections of vCA1, with a salt-and-pepper pattern observed within medial sections (**Fig. 1c-d**). We found that the order of tagging, tagging shock and then female exposure, did not impact the anatomical recruitment of these cells (**Extended Data Fig. 2**)

Next, using the same dual tagging strategy as described above, we asked whether or not these appetitive- and aversive-tagged populations were recruited during subsequent experiences of similar valences. To that end, we added a third time point by visualizing endogenous cFos expression 90 minutes after a final behavioral experience, i.e. exposure to sweetened condensed milk or restraint stress^19^ (**Fig. 1e-f**). To control the order of experiences, we counterbalanced the four groups, each of which contained two tagged populations of cells (i.e. “aversive” for cells tagged by fear conditioning and “appetitive” for cells tagged by male-female interactions), followed by a third experience of varying valence, restraint stress as another aversive and sweetened condense milk as the other appetitive experience, which was captured by endogenous cFos expression. Our groups were as follows: Aversive-Appetitive-Restraint Stress (GROUP 1), Aversive-Aversive-Restraint Stress (GROUP 2), Appetitive-Aversive-Sweetened Condensed Milk (GROUP3), and Appetitive-Appetitive-Sweetened Condensed Milk (GROUP 4) as shown in **Fig. 1f**. In each group we measured the cellular co-localization of TdTomato, EYFP, and cFos to infer which cells were active in one or more of the three experiences (**Fig. 1g**). All mice exhibited a similar proportion of tagged cells across all three tagging approaches regardless of method of tagging or valence (**Fig. 1h**, and **Extended Data Fig. 3**).

**Figure 2.**
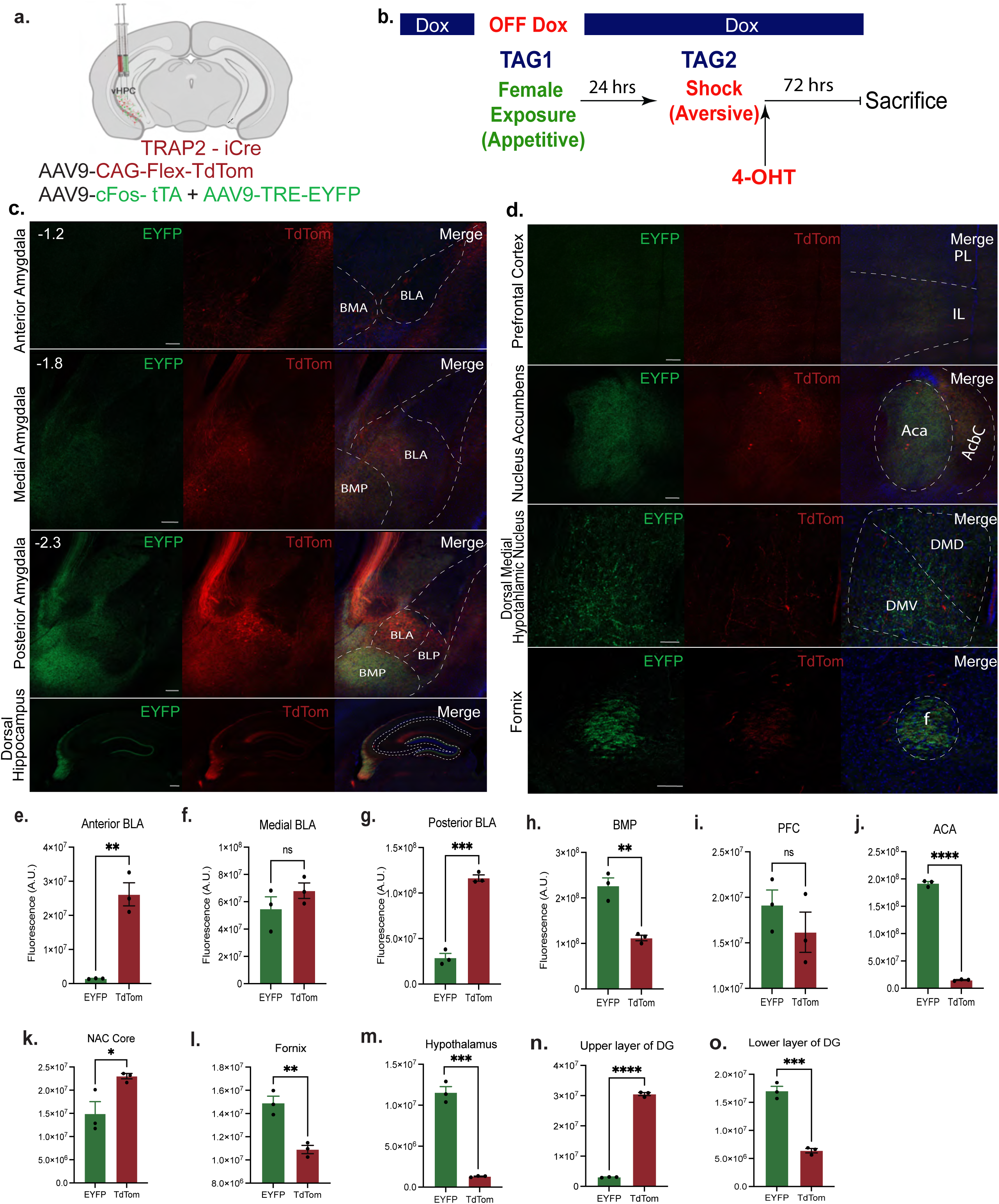
Ventral hippocampal neurons encoding appetitive or aversive valences have discrete projection patterns. **a,**Fos-CreERT2 mice were injected with a viral cocktail of 1:1 ratio of AAV9-c-Fos-tTA and AAV9-TRE-EYFP mixed in with AAV-Flex-DIO-TdTomato into the vHPC for dual memory tagging. **b,** Experimental timeline for tagging; EYFP represents aversive and TdTomato represents appetitive. **c-d,** Representative images of terminal projections; anterior, medial, and posterior BLA, dorsal hippocampus, prefrontal cortex, nucleus accumbens, dorsal medial hypothalamic nucleus, and the fornix. Arbitrary units for EYFP (appetitive) and TdTom (aversive) (N=3) for the **e,** anterior BLA (unpaired student’s t-test t=7.279, df=4, p= 0.00019)**, f,** medial BLA (t=1.271, df=4; p= 0.2725)**, g,** posterior BLA (t=14.95, df=4; p=0.0001) **, h,** BMP ( t=6.168, df=4; p=0.0035)**, i,** PFC (includes IL and PL) (t=1.078, df=4; p=0.3419), **j,** ACA (t=50.68, df=4; P<0.0001)**, k,** NAc Core (t=3.018, df=4; p=0.0392) **, l,** fornix (t=5.718, df=4; p=0.0046) **, m,** hypothalamus t=13.96, df=4; p=0.0002**, n,** upper layer of DG(t=54.94, df=4; p<0.0001)**, o,** lower layer of DG (t=11.73, df=4; p=0.0003).

Histological analyses revealed that there were significantly higher rates of cells showing co-localization of TdTomato and EYFP when labelled with the same experiences (i.e. appetitive and appetitive) and lower rates of co-localization when labeling differently valenced experiences (i.e. appetitive and aversive) in **Fig. 1i**. Further, we observed significant colocalization between TdTomato, EYFP, and cFos when mice were subjected to three aversive or three appetitive experiences (**Fig. 1l**), i.e. GROUP 2 and GROUP 4 respectively. Appetitive-tagged cells were preferentially reactivated when mice were exposed to sweetened condensed milk, and aversive-tagged cells were preferentially reactivated when mice were exposed to restraint stress (**Fig. 1j-k**). Together, these findings raise the intriguing possibility that vCA1 designates emotionally-relevant information to two partially non-overlapping sets of cells.

Moving forward, we chose two valenced engrams, shock and male-female exposure, as a proxy to better understand how the vHPC processes these aversive and appetitive experiences. Therefore, we used the same dual-tagging strategies to characterize the basic physiological properties of vHPC cells (**Extended Data Fig. 4**). TRAP2 mice were co-injected with the same activity dependent viruses and tagged in the same manner as mentioned previously (**Extended Data Fig. 4a-b**). Interestingly, we did not observe any differences in firing frequency, suprathreshold characterization, adaptation rate, input rate, or spike rates (**Extended Data Fig. 4c-l**) suggesting that, despite recruiting partially non-overlapping sets of cells for appetitive and aversive experiences, these tagged cells themselves share similar physiological characteristics.

Despite these physiological similarities, vHPC cells have been shown to project to a myriad of distinct brain regions involved in stress and approach-avoidance behaviors, thereby forming multi-regional networks involving emotion and memory^1^. We speculated that within these networks there exists structural heterogeneity partly defined by whether an experience is appetitively or aversely valenced, as has been observed in areas including the amygdala^10, 25, 27^, nucleus accumbens^29^, and medial prefrontal cortex^36^. Using our dual tagging strategy, we next traced vHPC outputs tagged by appetitive and shock experiences in a within-subject manner and measured the axonal fluorescence intensities in the following target areas, given their crucial role in emotional processing: BLA, NAc, PFC, dorsal hypothalamus, fornix, and the dentate gyrus (**Fig. 2a-b**). Interestingly, we observed both red and green fluorescent signal between appetitive and aversive-tagged vCA1 terminals in the medial BLA (A.P. -1.8) and the PFC (including IL and PL), while also finding evidence of structural segregation in the following regions: we observed predominantly stronger EYFP (appetitive) projections, as measured by fluorescence intensities, to the basomedial amygdala Posterior part (BMP), anterior commissure anterior part (ACA), fornix, dorsal medial hypothalamic nucleus (DMD and DMV), and the lower layer of the dentate gyrus (**Fig. 2c-o**). Further, we found predominantly stronger TdTomato (aversive) projections to the anterior BLA, posterior BLA, NAc core, and the upper layer of the DG (**Fig. 2c-o**). Previous studies have demonstrated that neurons in regions such as the PFC^36^, NAc^29^, and BLA^10.25^, collectively process experiences in a manner that can be anatomically segregated or heterogeneous. We posit that appetitive- and aversive-tagged vHPC terminals are embedded in a larger network of emotional memory processing that can be partly defined both by unique anatomical patterns and by the activity-dependent recruitment of ensembles involved in processing a specific valence.

Recent studies have identified unique molecular profiles of vCA1 cells containing distinct projection targets. These vCA1 projections transmit information to multiple areas involved in emotion and memory and form a network that can be organized by unique architectural features, including vCA1 cell inputs, outputs, and transcriptional signatures^1^. We wanted to ask the question of whether cell populations as a whole, for appetitive or aversive engram, have distinct genetic differences based on experience. Accordingly, we next examined whether or not the molecular composition of vHPC cells contained distinct genetic profiles. To catalogue the molecular landscape of vHPC cells in an activity-dependent manner, we tagged an appetitive experience (i.e. male-female interactions), a aversive experience (i.e. multiple shocks), or a neutral experience (i.e. exposure to the same conditioning cage without an appetitive or aversive stimuli; see methods; **Fig. 3a**). We then performed RNA-seq using tagged nuclei (EYFP+) isolated by Fluorescence-Activated Cell Sorting (FACS), which showed 0.5∼0.6% tagged nuclei from each group (**Fig. 3b-c**). Comparing aversive and appetitive vCA1 cells to the neutral group, we identified 474 differentially expressed genes (DEGs) in aversive vCA1 cells (**Fig. 3d** and **3f**), including 340 down regulated genes and 134 upregulated genes (**Fig. 3g**). We also identified 1,104 DEGs in appetitive vCA1 cells compared to the neutral group (**Fig. 3e** and **3f**), including 1,025 down regulated genes and 79 upregulated genes (**Fig. 3g**). There were 842 unique DEGs in appetitive engram cells and 212 unique DEGs in aversive engram cells (**Fig. 3f**), suggesting distinct transcriptional landscapes in these two populations.

**Figure 3.**
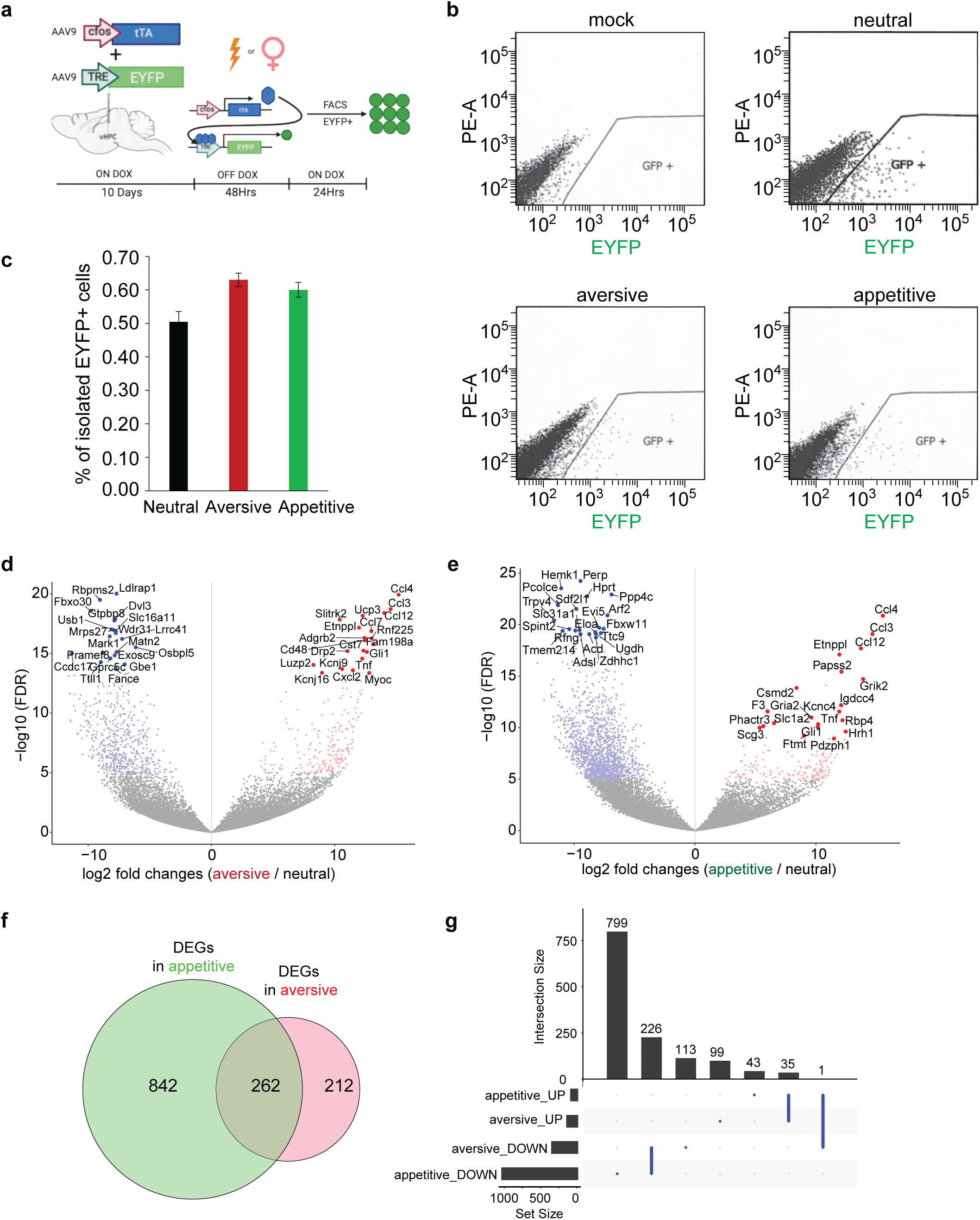
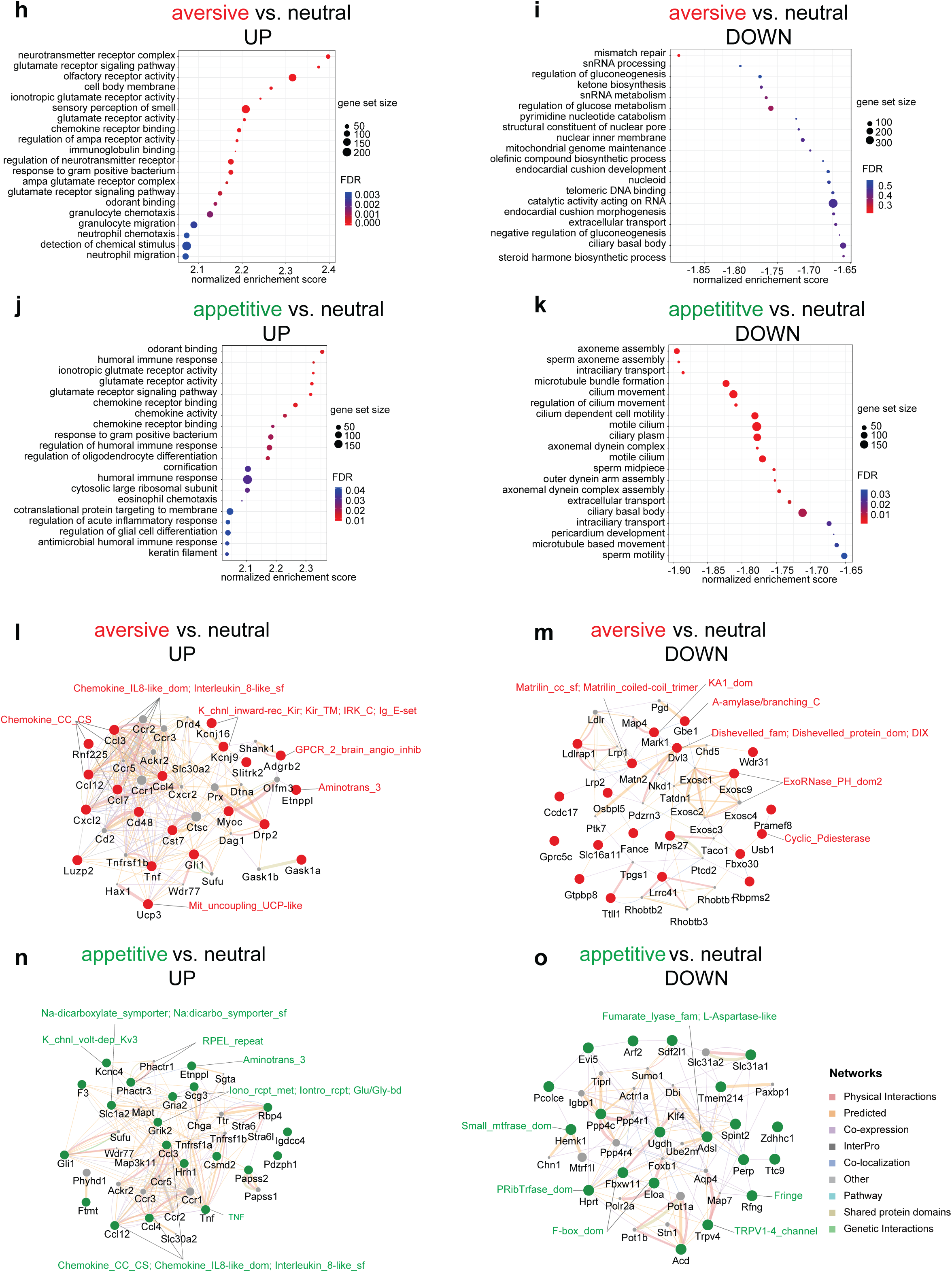
Distinct molecular landscapes of appetitive and aversive hippocampus engram cells. **a**, Experimental scheme to label, isolate, and analyze the two groups of vCA1 cells labeled by EYFP upon aversive stimulation (electrical shock) or appetitive stimulation (male mice engaged in social interaction with female mice) or neutral conditioning in the same cage without aversive or appetitive stimulation. **b.** FACS isolation of EYFP-appetitive nuclei from the hippocampi of mice labeled by AAV9-cFos-tTA + AAV9-TRE-ChR2-EYFP virus from their respective groups **c**, Percentage of EYFP-appetitive nuclei from each group. **d**, A Volcano plot with the relative fold change of gene expression in log 2 ratio and the FDR-adjusted p-value in log 2 ratio as X and Y axis showing the up and down regulated genes in aversive vCA1 cells compared to neutral group. Up or down regulated genes with FDR adjusted p-value less than 1e-5 plus at least four-fold differences were highlighted in either red or blue respectively. The gene names were displayed for the top 20 most significant genes as sorted by Wald statistics. **e**, A Volcano plot showing the up and down regulated genes in appetitive vCA1 cells compared to neutral group. **f**, A Venn diagram showing the 1,104 differentially expressed genes (DEGs) in appetitive and 474 DEGs in aversive vCA1 cells. 262 DEGs were identified in both groups. **g**, An UpSet plot showing 226 down regulated genes in both aversive and appetitive vCA1 cells, and 35 up regulated genes in both aversive and appetitive vCA1 cells. Only one gene (*Nufip1*) was upregulated in appetitive engram but down regulated in aversive engram. **h**, Gene Enrichment Set Analysis of the up regulated pathways in aversive engrams using Gene Ontology (GO) module. The size of the dot represents the gene count and the color of the dot indicates the FDR. The pathway with an FDR-adjusted P value smaller than 0.25 is considered as significantly enriched. **i**, GO analysis of down regulated pathways in aversive vCA1 cells. **j**, GO analysis of up regulated pathways in appetitive vCA1 cells. **k**, GO analysis of down regulated pathways in appetitive vCA1 cells. GeneMANIA network of the top 20 upregulated and downregulated genes in aversive and appetitive vCA1 cells, respectively. Genes with red nodes are upregulated **l**, or downregulated **m**, in the aversive vCA1 cells compared to the neutral sample, and genes with green nodes are upregulated (**n**) or downregulated (**o**) in the appetitive vCA1 compared to the neutral sample. The grey nodes represent the genes interacting with these 20 differentially expressed genes. The shared protein domains supported by InterPro domain databases within each network were labeled in red for aversive vCA1 cells and in green for appetitive vCA1 cells. The detailed description for each shared protein domain is listed in Table S4. The interaction network categories between these genes are annotated in the legend next to panel **o**.

Furthermore, the top 20 downregulated DEGs identified for aversive and appetitive vCA1 cells showed no overlap with each other, and 14 among the top 20 upregulated DEGs for aversive and appetitive vCA1 cells do not overlap with each other (**Fig. 3d** and **3e**, **Table S1** and **S2**), supporting the distinct transcriptomes in these neurons. Interestingly, among the 262 shared DEGs we identified one gene ***Nufip1*** (Nuclear Fragile X Mental Retardation Protein Interacting Protein 1) that was upregulated in appetitive vCA1 cells and downregulated in aversive vCA1 cells (**Fig. 3g**). This gene encodes a nuclear RNA binding protein that contains a C2H2 zinc finger motif and a nuclear localization signal^47^. Diseases associated with *NUFIP1* mutations include Peho Syndrome (progressive encephalopathy with Edema, Hypsarrhythmia and Optic atrophy), an autosomal recessive and dominate, progressive neurodegenerative disorder that starts in the first few weeks or months of life. Its interacting protein FMRP1 is essential for protein synthesis in the synapse^48^ and CGG trinucleotide expansion mutation of FMR1 gene coding FMRP1 cause Fragile X syndrome, the most common intellectual disability in males^49^. Further investigation of the functional significance of NUFIP1 and other DEGs could reveal mechanistic insights to the transcriptomic plasticity engaged by the valences of memory.

To gain deeper insight into the molecular signatures of transcriptomes associated with aversive and appetitive vCA1 cells, we performed Gene Ontology (GO) analysis of the up- and down-regulated pathways in aversive and appetitive vCA1 cells (**Fig. 3h-k**). We found that the top upregulated pathway in aversive vCA1 cells involved neurotransmitter complexes such as ionotropic glutamate receptor activity (**Fig. 3h**). This finding is consistent with previous studies using brain tissues that identified 3,759 differentially methylated DNA regions in the hippocampus associated with 1,206 genes enriched in the categories of ion gated channel activity after contextual fear conditioning^48, 50, 51^. Interestingly, we found the top downregulated pathway in aversive vCA1 cells involved DNA mismatch repair (**Fig. 3i**). Although the top upregulated pathways in appetitive vCA1 cells also include ionotropic glutamate receptor activity (**Fig. 3j**), the top downregulated pathways in appetitive vCA1 cells enrich on axoneme assembly and microtubule bundle formation (**Fig. 3k**) different from the pathways downregulated in appetitive vCA1 cells (**Fig. 3i**). Next, we compared the RNA-seq data between aversive and appetitive vCA1 cells directly. We identified 494 DEGs, including 47 upregulated genes and 447 downregulated genes (**Extended** **Fig. 5a**, **Table S3**). Furthermore, we found that five pathways were upregulated in appetitive vCA1 cells such as nuclear exosome RNAse complex and 16 pathways were down regulated in appetitive engram compared to aversive vCA1 cells, such as axoneme assembly (**Extended Fig. 5b** and **5c**). These differentially altered signaling pathways between aversive and appetitive vCA1 cells, which resulted from multiple comparisons, support our conclusion that these neurons indeed represent transcriptionally distinct subpopulations. One future direction is to explore the functions of these DEGs and altered signaling pathways. As a proof-of-concept, we applied GeneMANIA^54^ to predict the functions of the top 20 DEGs in **Fig. 3l-o**. For instance, the brain-specific angiogenesis inhibitor 2 (BAI2) is uniquely identified for the DEGs upregulated in aversive vCA1 cells (**Fig. 3l**), and the Ionotropic glutamate receptor is only implicated for the DEGs upregulated in appetitive vCA1 cells (**Fig. 3n**). Similarly, the WNT signaling pathway component Dishevelled is uniquely identified for the DEGs downregulated in aversive vCA1 cells (**Fig. 3m**), and the TRPV1-4 channel is only implicated for the DEGs upregulated in appetitive vCA1 cells (**Fig. 3o**). Loss- and gain-function study of these predicted genes will provide mechanistic insight for the molecular signatures of engram neurons with different memory valences.

Besides the distinct transcriptomic profiles in engram neurons described above, recent studies reveal the transcriptional priming role of epigenetic regulation in engram^43^. To explore whether the dynamic epigenomic landscape also contributes to the specificity of engram neurons with different valences, we performed reduced representation bisulfite sequencing (RRBS) to characterize the DNA methylation landscapes in aversive and appetitive vCA1 cells. As shown in **Fig. 4a** and **4b**, we identified 1939 differentially methylated cytosines (DMCs) with the change of DNA methylation larger than 20% and p value smaller than 0.05 in aversive vCA1 cells compared to the neutral group, and 3117 DMCs in appetitive vCA1 cells. These DMCs are located in the different positions of the genome including 5’UTR, promotor, exon, intron, 3’UTR, transcription termination sites (TTS), intergenic, non-coding regions, suggesting different functional output at the transcriptional level. Interestingly, the genomic distributions of these DMCs in aversive vCA1 cells are slightly different from the one in appetitive vCA1 (**Table S5**). Based on these DMCs, we identified differentially methylated regions (DMRs) that contain at least two DMCs for each DMR (**Fig. 4c** and **4d**). These DMRs allow us to identify the differentially methylated genes (DMGs) that either contain or close to these DMRs. The top 20 DMGs with the change of methylation level larger than 20% and p value smaller than 0.05 show no overlapping between aversive and appetitive vCA1 cells, suggesting different memory valences trigger different changes of DNA methylations. Among the 266 DMGs in appetitive vCA1 cells and the 98 DMGs in aversive vCA1 cells, only 32 DMGs are commonly shared (**Fig. 4e**, **Table S6**), confirming the distinct DNA methylation landscape between these two populations of engram cells. Last, we performed Gene Ontology (GO) analysis of DMGs in aversive and appetitive vCA1 cells (**Fig. 4f** and **4g**). We found that the pathways in aversive vCA1 cells mainly enriched in the structure and function of synaptic connections (**Fig. 4f**). However, the enriched pathways in appetitive vCA1 cells are much more diverse including axon growth, synaptic connection, ion channels, and RNA polymerase II transcription regulator complex (**Fig. 4g**). These differentially enriched pathways between aversive and appetitive vCA1 cells suggest potentially distinguished functional outputs attributed by DNA methylation at the transcriptional level to confer the specificity of memory valences. One interesting future direction is to explore the maintenance and functions of these DNA methylation changes during the consolidation and recall of memory. Overall, our results in **Fig. 3** and **4** showed that the distinct molecular signatures of aversive and appetitive vCA1 cells are reflected at the transcriptomic and epigenomic levels likely contributing to the different valences of memory.

**Figure 4.**
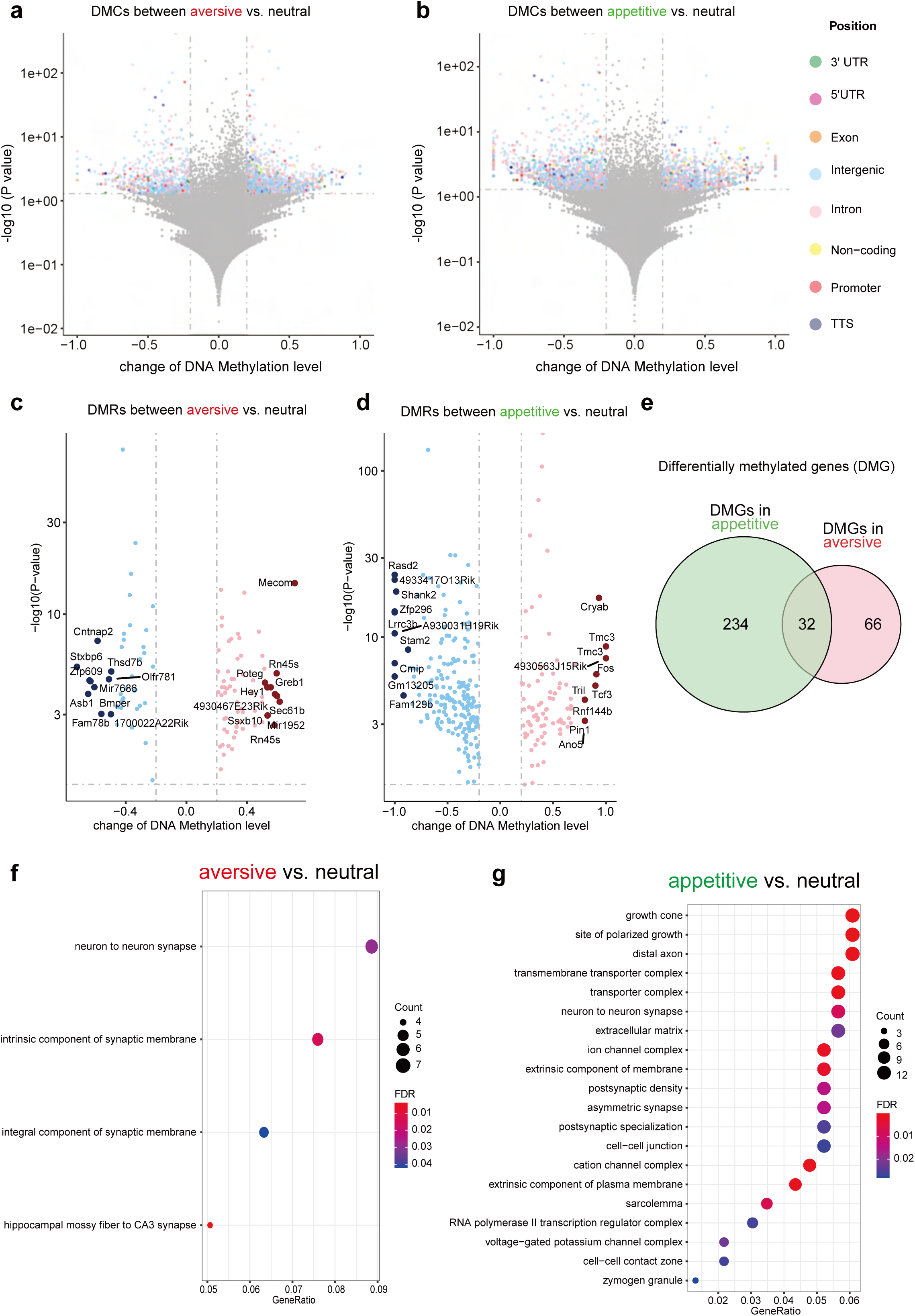
Distinct DNA methylation landscapes of aversive and appetitive vCA1 cells. Volcano plots showing the differentially methylated cytosines (DMC) with the change of DNA methylation larger than 20% and p value smaller than 0.05 in aversive **a,** or appetitive **b,** vCA1 cells compared to the neutral group. DMCs in different genomic positions (3’UTR, 5’UTR, exon, intergenic, intron, non-coding promotor, TSS) are color coded. Volcano plot showing differently methylated regions (DMRs) that contain at least two DMCs for each DMR in aversive **c**, and appetitive **d**, vCA1 cells. The top 10 hypermethylated and hypomethylated regions were highlighted in either pink or blue separately and were annotated to its nearest genes. **e,** A Venn diagram showing differently methylated genes (DMGs) that are associated with DMRs in c and d identified in aversive and appetitive vCA1 cells. **f**, GO analysis of DMGs in aversive vCA1 cells. **g**, GO analysis of DMGs in appetitive vCA1 cells.

vCA1 is known to have monosynaptic projections to the BLA, NAc, and the mPFC^24^. Previous studies have shown that these brain regions are important in the modulation of both appetitive and aversive experiences, especially in the NAc^29, 34^ and the BLA^25, 26^, in which molecularly and topographically distinct cellular populations have been identified for each behavior. Therefore, we tested for a causal role of tagged vHPC cell bodies and its selected terminals in driving behavior by first infusing a virus cocktail of AAV9-cFos-tTA and AAV9-TRE-ChR2-EYFP or AAV9-TRE-EYFP into vCA1. We then implanted an optical fiber bilaterally above either the vCA1 or its terminals, vCA1–BLA, vCA1–NAc, vCA1–PFC (**Fig. 5a**). We first found that all terminals were capable of activating their corresponding downstream targets by assessing increases in cFos levels following stimulation (**Extended Data Fig. 6**). Following 10 days of recovery, a separate group of mice were taken off DOX and tagged with aversive (i.e. shock) or appetitive (i.e. male-female interactions) experiences. As illustrated in **Fig. 5b**, for the first set of experiments, on Day 1, the aversively tagged mice were placed in a real-time place preference/avoidance (RTPP/A) chamber on day 1 to assess baseline levels. On day 3, the animals were placed back into the PTPP/A chamber; this time they received optical stimulation at 20Hz bilaterally on one side and no stimulation on the other. We found that optical stimulation of vCA1–BLA or vCA1–NAc terminals drove aversion (**Fig. 5e & f**), whereas the EYFP controls, vCA1 cell body stimulation, and vCA1-PFC terminals did not statistically deviate from baseline levels (**Fig. 5c, d, & g**). On day 5 of the experiment, the mice were subjected to an induction protocol, as previously reported^13^, to test the capacity of the engram to “switch” the behavior it drives. The aversive-tagged male mice were placed in a new chamber with a female mouse for 10 minutes while receiving optical stimulation for the entire duration of exposure. Afterwards, the animals were placed back in their chambers and assessed for behavioral changes on day 7. In this post-induction test, we observed that optical stimulation of vCA1–BLA terminals was now sufficient to drive preference despite driving aversion in the pre-induction test earlier (**Fig. 5e**). The induction protocol also revealed that vCA1–NAc terminals, which previously were sufficient to drive aversion, now had reversed or “reset” their capacity to modulate behavior and returned to baseline levels (**Fig. 5g**). Lastly, we saw no changes in the EYFP controls, vCA1 cell body, or vCA1–PFC stimulation (**Fig. 5c, d, & g**).

**Figure 5.**
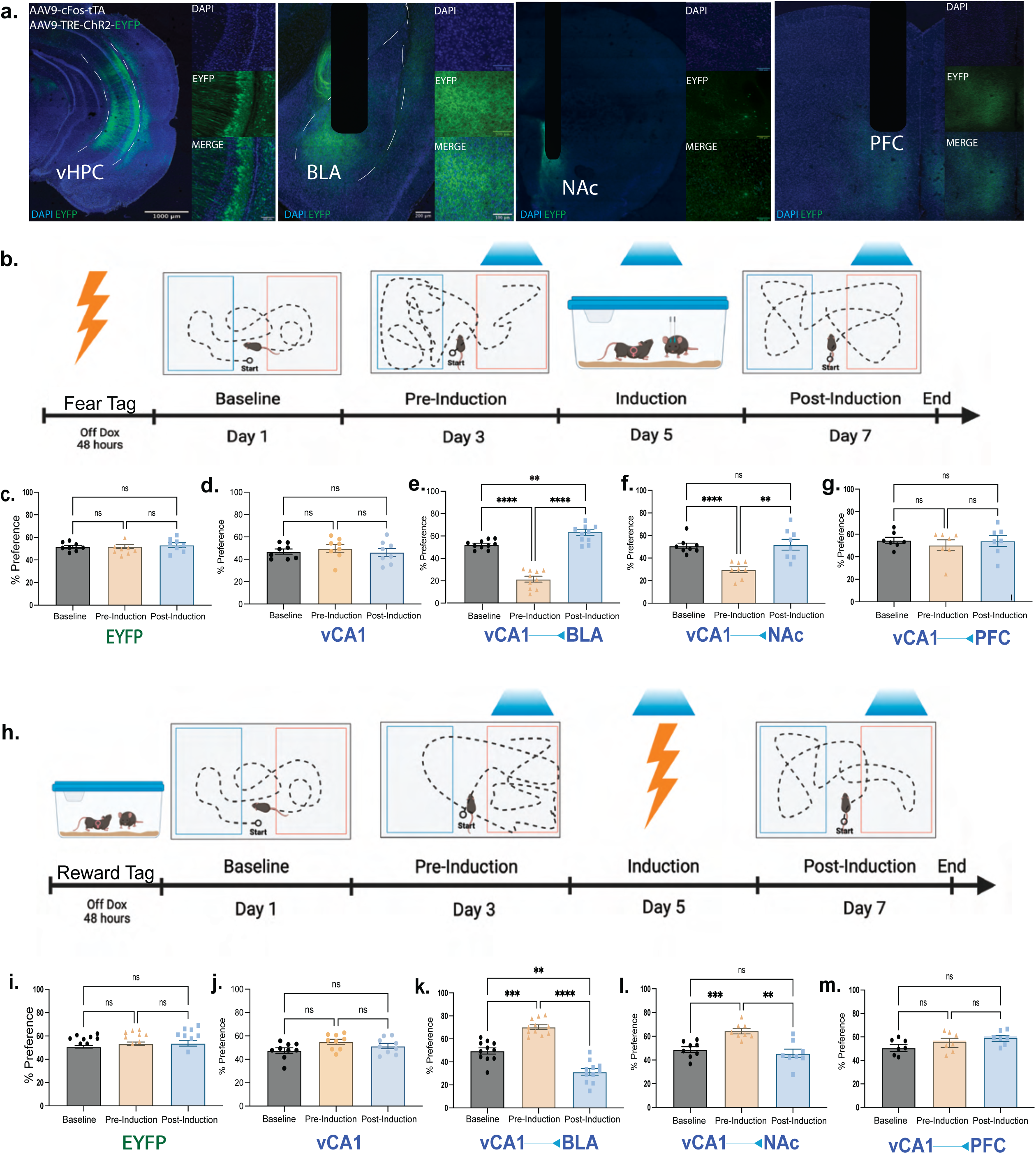
Hippocampal valence specific outputs are sufficient to drive preference or aversion in a projection-specific and functionally reversible manner. Mice were injected with a virus cocktail of AAV9-c-Fos-tTA AAV9-TRE-ChR2-EYFP or AAV9-TRE-EYFP into vCA1 and optic fibers were placed bilaterally over the cell bodies of vCA1 or the terminals from vCA1 to the BLA, Nacc, or the PFC, in separate groups. The mice were then tagged with either a aversive or appetitive experience and subjected to real time opto-place avoidance and preference. **a**, Representative images of ChR2-EYFP labeling cell bodies in vCA1 and its respective terminals in the BLA, NAcc, and PFC. **b**, Fear to reward protocol in which the subjects received stimulation of vCA1 cell body or vCA1 projections to the BLA, NAcc, or PFC terminals. **c-g**, Percent preference for the stimulation side at baseline, pre-induction, and post-induction (n = 7 subjects for EYFP controls, n=8 subjects for vCA1, n = 8 subjects for BLA, n = 7 for NAcc, and n = 7 for PFC, **P = 0.0018, ***P = 0.0006, repeated measures one-way ANOVA followed by Tukey’s multiple comparison test). **h**, Reward to Fear protocol in which the subjects received stimulation of BLA, NAcc, or PFC terminals originating from vCA1. **i-m**, Percent preference for the stimulation side at baseline, pre-induction, and post-induction (n = 7 for EYFP, n=9 for vCA1, n = 8 for BLA, n = 7 for NAcc, and n = 7 for PFC, **P = 0.0032, ****P < 0.0001, repeated measures one-way ANOVA followed by Tukey’s multiple comparison test).

Next, we asked whether appetitive-tagged vCA1 terminals to the BLA, NAc, or PFC were sufficient to modulate behavior (**Fig. 5h**). Similar to the above findings, optical stimulation of vCA1–BLA (**Fig. 5k**) and vCA1-NAc (**Fig. 5l**) terminals, but not in any of the other groups (**Fig. 5i, j, & m**), was sufficient to drive place preference. During the induction phase of the experiment, we placed the mice into a fear conditioning chamber where they received multiple foot shocks and simultaneous optical stimulation of the appetitive-tagged terminals. This experiment assessed if these vCA1 terminals are able to “switch” or “reset” their capacity to drive preference in a manner mirroring the experiments above. Indeed, we found that in the post-induction tests, the vCA1–BLA group switches from driving preference to aversion (**Fig. 5k**) while the vCA1–NAc group resets from driving preference back to baseline levels (**Fig. 5l**). Furthermore, we did not observe this effect in neutral-tagged animals (**Extended Data Fig. 7**), nor did we see any statistically significant behavioral changes for the RTPP/A task in the EYFP controls, vCA1 cell body stimulation group, or vCA1–PFC stimulation group (**Fig. 5i, j, & m**). Importantly, we tested whether optical stimulation may cause increases in motor behaviors. We found no significant difference in distance travelled across all groups during light on or off epochs in an open field (**Extended Data Fig. 8**). Moreover, the lack of preference or avoidance observed from vCA1 cell body stimulation raised the possibility that vCA1’s role in driving behavior is determined in a terminal-specific manner^24^; a notion that dovetails with recent studies suggesting that computations along the axons of a given cell body can differentially drive behavior in accordance with the downstream target^8^. Taken together, these results suggest that vHPC cell bodies relay their behaviorally-relevant and valence specific content to its downstream targets despite sharing partially non-overlapping molecular and neuronal signatures and distinct projection patterns. This is corroborated by evidence showing that vHPC axonal outputs preferentially route independent features of a given behavior^7^.

Here we have shown that the vHPC processes appetitive and aversive experiences in defined populations of cells that are partially distinct at the molecular, cellular, and projection-specific levels. We also demonstrated their capacity to drive behaviors through functionally plastic projection-specific terminals. Our immunohistochemical data suggest that vCA1 contains at least three populations of neurons: two subsets that can be demarcated based on their anterior-posterior locations and preferential response to appetitive or aversive valences, and a third population that responds to both, perhaps reflecting a biological predilection for salience.^5^ While their exact brain-wide structural and functional outputs remain undetermined, we speculate that our observed population of vCA1 cells responding to aversion are a superset of recently observed anxiety cells that transmit information to the hypothalamus^27^ and PFC^28^. Moreover, vCA1 cells processing aversive or appetitive perhaps route information to and innervate the BLA and NAc at differing anatomic, receptor- and cell-type specifically optimizing their capacity to integrate mnemonic information^27^.

Additionally, by combining transgenic activity-dependent tagging strategies with all-virus-based expression of fluorophores, our design permits the visualization of multiple ensembles in a within-subject manner, which coalesces with previous studies monitoring and manipulating a single ensemble active at two discrete time points as well^29^. By intersecting these approaches with genetic sequencing strategies, these tools enable the tagging, manipulation, and molecular documentation of cells processing aversive and appetitive behaviors opening the possibility of cataloguing topographical similarities and differences between the two in a brain-wide manner. For instance, future studies may probe the molecular composition of appetitive- and aversive-tagged vCA1 cells along its anterior-posterior axis, and test whether or not the transcriptomic profiles of these cells differ both across valence and their physical location. Moreover, it is intriguing that vCA1 appetitive- and aversive-tagged terminals showed evidence of segregation within the amygdala and hippocampus. Consequently, subsequent research may seek to functionally tease out their contributions to behavior, ands measure the types of information they are transmitting through a combination of imaging and terminal-specific perturbation approaches.

It is important to constrain interpretations of engrams given the vast array of genetic tools used to access cells in an activity-dependent manner. For instance, vCA1 ensembles vary drastically in size and in activity patterns across learning and memory. These numbers range from single percentages and can reach ∼35% of cells depending on immediate early gene marker used, tagging strategy, rodent line, and viral tools employed^32–35^. Fittingly, we believe that our dual-tagging strategy partly over-samples the number of tagged cells given the time-frame necessary for tagging to occur (e.g. 48 hours off Dox; 72 hours post-4OHT injection^37–40^), and future experiments may aim to improve the temporal resolution of contemporary tagging approaches to enhance the signal-to-noise ratio of experience-related tagging to background tagging or leakiness. Additionally, cFos+ cells only reflect a subset of an engram that is otherwise distributed throughout the brain and recruits numerous immediate early genes, cell types, and complementary physiological activity that activity-dependent markers may fail to capture due to the relatively slow timescales of gene expression. We indeed note that while cFos+ cells indicate recent neural activity, a cell that was recently active does not necessarily have to express cFos especially across brain regions and cell types which presumably have their own thresholds for cFos expression. These points highlight the complex nature of a memory engram and underscore a caution that is warranted when interpreting data in light of inherent technical limitations. We believe that a multi-faceted approach combining genetic tagging strategies with real-time imaging during complex behaviors will help to disentangle the relationship between neural activity, genetic modifications, and systems-level changes in response to learning and memory.

Our terminal manipulations are in line with recent studies demonstrating terminal-specific routing of information from vCA1 cell bodies through a variety of single, bi-, and trifurcating processes^8, 41, 42^. Our data provide a “gain-of-function” demonstration that activated vCA1 terminals can drive preference or aversion, which we believe was obfuscated by cell body stimulation given that the latter presumably activates the cell body and most, if not all, corresponding axonal outputs. The former would selectively modulate a set of terminals emerging from vCA1 that project to a distinct target area while only minimally affecting vCA1 terminals projecting to other target areas. Additionally, while the molecular basis underlying our valence “switch” experiments remain unknown, we posit that the plasticity of the transcriptome could confer the ability to the switch which aspect of behavior a terminal drives given the rapid and enduring responses to learning and memory present at the genetic level in tagged cells.^43^

Indeed, the comprehensive transcriptional landscape in the mouse hippocampus is dynamic across the lifespan of memory formation and recall, and its experience-dependent modifications are largely present specifically in tagged cells within minutes of learning and lasting for several weeks^42^. Future experiments may perform RNA-seq on cells before and after the induction protocol to measure the ensuing transcriptional changes and identify putative loci mediating such functional plasticity. Further, our sequencing data enhances our understanding of how the ventral hippocampus genetically parses out both appetitive and aversive engrams, how those experiences can cause a change in the upregulation and down-regulation of discrete genes, and how these experiences can have lasting effects on the epigenetic genome through the methylation study (**Fig. 4**). Future studies, for instance, can build on recent work assessing the molecular and projection-defined connectivity between vCA1 and its downstream targets with MAPSeq^1^, while adding an activity-dependent component, such as our dual memory tagging approach, to measure organizing principles of the ventral hippocampus and its projections in a manner that takes a cell’s history into account. This study in particular provides an influential platform for characterizing the organization of the ventral hippocampus and its non-random input-output patterns, and we posit that this logic includes an activity-dependent-defined dimension such that appetitive and aversive experiences engage unique input-output hippocampal circuitry as well.

Finally, while the physiological basis by which terminals can switch or reset their capacity to drive valence-specific behaviors remains unclear, future studies may consider candidate mechanisms including homeostatic plasticity, dendritic growth and retraction, and counter-conditioning-facilitated changes along the axon-dendrite interface between vCA1 and the BLA or NAc^25, 26^. In line with this speculation, previous studies have demonstrated that the dorsal hippocampus contains defined sets of functionally plastic cell bodies sufficient to drive aversive or appetitive behavior, while the BLA contains fixed populations that are sufficient to drive each behavior contingent on the anatomical location stimulated along the anterior-posterior axis as well as on their projection-specific sites.^7, 8, 25–27^ Subsequent research can be aimed at utilizing our dual-memory tagging approach to genetically and physiologically map out which cell-types and circuits show such hardwired or experience-dependent responses to emotional memories as well. In our study, we speculate that vCA1 cells become appetitive or aversive in an experience-dependent manner as opposed to being hardwired for either valence per se, as has been observed in many other brain regions (e.g. BLA^25^). However, though it remains possible that a subset of these vCA1 cells are preferentially tuned to process a given valence in an experience-independent manner.

Indeed, the notion that experience itself modifies these neurons to become appetitive or aversive does not preclude the possibility that subsequent experiences will modify their capacity in a flexible manner to drive a given behavior associated with a given valence.

Moreover, in our study, while the criteria for valence was met in **Fig. 1** (e.g. hippocampus cells responded differentially to stimuli of positive and negative valence in a manner independent of stimulus features), our subsequent data sets honed in on a single appetitive (e.g. male-female social interactions) and a single aversive (e.g. foot shocks) experience for thorough molecular, anatomical, and behavioral profiling. Importantly, we highlight here that in order to make a direct claim about valence, the task structure needs to be held constant. For instance, an alternative interpretation of our current data is that our observed differences in gene enrichment profiles and anatomical segregation are due to the inherent differences in tasks used (e.g. male-female social interactions vs. contextual fear conditioning), and not valence per se. It remains possible therefore that male-female social interactions, which require multi-modal integration of social and contextual cues, recruit a unique set of genes and ventral hippocampal anatomy in comparison to contextual fear conditioning, in which a conditioned cue (e.g. context) is associated with an unconditioned cue (e.g. shock), and therefore recruits task-specific molecular and anatomical activity. As such, we propose that future studies may focus on tagging experiences in which most, if not all, environmental features are held constant except the valence associated with a unimodal stimulus itself, which we believe opens up an experimental platform for studying how multiple experiences of similar or varying valence differentially engage cellular populations. While we believe that this partially accounted for by utilizing multiple experiences of similar valence in **Fig. 1**, additional future experiments may implement an in vivo recording approach in which putative negative- and positive-tagged cells are imaged while task structure and valence are parametrically varied. Building from this notion, our RNA-Seq and DNA methylation data provides an additional means by which memories alter the functions of the genome in a valence-dependent manner, both in healthy and pathological states. For instance, our methylation data identified Pin1^60, 61^ among 266 DMGs in the appetitive vCA1 cells, which is known for its neuroprotective qualities specifically in neurodegeneration through the regulation in the spread of cis p-tau. Follow-up studies may leverage these appetitive engrams by altering these DMGs as a means to alleviate psychiatric and neurodegenerative diseases.

Together, we propose that, in addition to processing spatio-temporal units of information, the hippocampus contains discrete sets of cells processing aversive and appetitive information that relay content-specific and behaviorally-relevant information to downstream areas in a molecularly-defined and projection-specific manner, thus collectively providing a multi-synaptic biological substrate for multiple memory engrams.

## ONLINE METHODS

### Subjects

Fos^CreER^ (Jax stock: #021882) and Wildtype male C57BL/6 mice (2-3 months of age; Charles River Labs) were housed in groups of 5 mice per cage. The animal vivarium was maintained on a 12:12-hour light cycle (lights on at 0700). Mice were placed on a diet containing 40 mg/kg doxycycline (Dox) for a minimum of 48 hours prior to surgery with access to food (doxycycline diet) and water *ad libitum*. Mice were given a minimum of ten days after surgery to recover. Dox-containing diet was replaced with standard mouse chow (*ad libitum*) 48 hours prior to behavioral tagging to open a time window of activity-dependent labeling^1, 2^. All subjects were treated in accordance with protocol 17-008 approved by the Institutional Animal Care and Use Committee at Boston University.

### Stereotaxic Surgery and Optic Implant

Stereotaxic injections and optical fiber implants follow methods previously reported.^1, 2^ All surgeries were performed under stereotaxic guidance and subsequent coordinates are given relative to bregma (in mm) dorsal ventral injections were calculated and zeroed out relative to the skull. Mice were placed into a stereotaxic frame (Kopf Instruments, Tujunga, CA, USA) and anesthetized with 3% isoflurane during induction and lowered to 1-2% to maintain anesthesia (oxygen L/min) throughout the surgery. Ophthalmic ointment was applied to both eyes to prevent corneal desiccation. Hair was removed with a hair removal cream and the surgical site was cleaned three times with ethanol and betadine. Following this, an incision was made to expose the skull. Bilateral craniotomies involved drilling windows through the skull above the injection sites using a 0.5 mm diameter drill bit. Coordinates were -3.16 anteroposterior (AP), ±3.1 mediolateral (ML), and -4.6 dorsoventral (DV) for vCA1; - 1.8 AP, ± 3.1 ML, and -4.7 DV for the BLA; -1.25 AP, ± 1.0 ML, and -4.7 DV for the NAc; 1.70 AP, ± 0.35 ML, and -2.8 DV for the PFC. All mice were injected with a volume of 300nl of cocktail per site at a control rate of 100 μl min^-1^ using a mineral oil-filled 33-gage beveled needle attached to a 10 μl Hamilton microsyringe (701LT; Hamilton) in a microsyringe pump (UMP3; WPI). The needle remained at the target site for five minutes post-injection before removal. For all targets, bilateral optic fibers were placed 0.5 DV above the injection site. Jewelry screws secured to the skull acted as anchors. Layers of adhesive cement (C&B Metabond) followed by dental cement (A-M Systems) were spread over the surgical site. Mice received 0.1 mL of 0.3 mg/ml buprenorphine (intraperitoneally) following surgery and were placed on a heating pad during surgery and recovery. Histological assessment verified viral targeting and fiber placements. Data from off-target injections were not included in analyses.

### Activity-dependent Viral Constructs

pAAV9-cFos-tTA, pAAV9-TRE-eYFP and pAAV9-TRE-mCherry were constructed as previously described (Ramirez et al., 2015). AAV9-c-Fos-tTA was combined with AAV9-TRE-eYFP or AAV9-TRE-ChR2-EYFP (UMass Vector core) prior to injection at a 1:1 ratio. This cocktail was further combined in a 1:1 ratio AAV9-CAG-Flex-DIO-TdTomato (UNC Vector Core).

### Optogenetic Methods

Optic fiber implants were plugged into a patch cord connected to a 450nm laser diode controlled by automated software (Doric Lenses). Laser output was tested at the beginning of every experiment to ensure that at least 15 mW of power was delivered at the end of the patch cord (Doric lenses).

### Behavior Tagging

When animals were off Dox, as previously reported^9–12^, Dox diet was replaced with standard lab chow (*ad libitum*) 48-hours prior to behavioral tagging. Female exposure: One female mouse (PD 30-40) was placed into a clean home cage with a clear cage top. The experimental male mouse was then placed into the chamber and allowed to interact freely for 2 hours. Fear exposure: Mice were placed into a conditioning chamber and received four 1.50mA (2s) foot shocks over an 8-minute training session. Following tagging, Dox was reintroduced to the diet and the male mice were returned to their home cage with access to Dox diet. For 4-OHT tagging, 4-hydroxytamoxifen (Sigma: H7904) was diluted into 100% ethanol and vortexed for 5 minutes. Once in solution, corn oil was added and the solution was sonicated to achieve a dilution of 10mg/kg stock. When the solution was ready, 4-OHT was loaded into syringes for injection and any left-over solution was placed in the -20C to be used in the future. On the day of tagging, 200mg of 10mg/ml stock (2mg 4OHT) was administered I.P. in Fos^creER^ mice 30 minutes following behavior and were left undisturbed for 72 hours. Mice were injected with saline or DMSO at least twice prior to tagging protocol to acclimate animals to injection and prevent off target tagging.

### Behavioral Assays

All behavior assays were conducted during the light cycle of the day (0700– 1900). Mice were handled for 3–5 days, 5-10 minutes per day, before all behavioral experiments. The testing chamber consisted of a custom-built rectangular box with a fiber optic holder (38 x 23.5 x 42 cm). Red tape divided the chamber down the middle, creating two halves. Right and left sides for stimulation were randomized. Day 1 was used to assess baseline levels, during which the mouse was given 10 minutes to freely explore the arena. Animals were rerun in baseline days if they showed more than a 45-55% preference for either side. Animals were acclimated to the chamber until they had a minimum of a 45/55 preference. Once baseline is achieved, the day following the first engram tag, mice received light stimulation (15 ms pulses at 20-Hz) upon entry in the designated side of the chamber (counterbalanced across groups) over a 10-minute test period. Once the mouse entered the stimulated side, a TTL signal from the EthoVision software via a Noldus USB-IO Box triggered a stimulus generator (STG-4008, Multi-channel Systems). 6A video camera (Activeon CX LCD Action Camera) recorded each session and an experimenter blind to treatment conditions scored the amount of time on each side. Statistical analyses involved one-way ANOVAs comparing group difference scores [time (in seconds) on stimulated side minus time on unstimulated side] as well as changes across days using matching or repeated measures one-way ANOVA. Behavioral diagrams were made with BioRender.

### Optogenetic Induction Protocol

For optogenetic **fear induction**, subjects were placed in a shock chamber with light stimulation (20 Hz, 15ms) for 500 seconds. Foot shocks (1.5 mA, 2s duration) were administered at the 198s, 278s, 358s, and 438s time points. During optogenetic **reward induction**, in a different room from the initial fear tag, the subject was placed into a clean homecage with a female mouse. Light stimulation (20Hz, 15ms) was applied to the male mouse for 10 minutes.

### Immunohistochemistry

Immunohistochemistry follows protocols previously reported^15, 16, 18^. Mice were overdosed with 3% isoflurane and perfused transcardially with cold (4° C) phosphate-buffered saline (PBS) followed by 4% paraformaldehyde (PFA) in PBS. Brains were extracted and stored overnight in PFA at 4°C. Fifty μm coronal sections were collected in serial order using a vibratome and collected in cold PBS. Sections were blocked for 1 hour at room temperature in PBST and 5% normal goat serum (NGS) or Bovine albumin serum (BSA) on a shaker. Sections were transferred to wells containing primary antibodies (1:1000 guinea anti-c-Fos [SySy]; 1:1000 rabbit anti-RFP [Rockland]; 1:5000 chicken anti-GFP [Invitrogen]) and allowed to incubate on a shaker overnight at room temperature or 3 days at 4 degrees C. Sections were then washed in PBST for 10-min (x3), followed by 2-hour incubation with secondary antibody (1:200 Alexa 555 anti-rabbit [Invitrogen]; 1:200 Alexa anti-guinea 647 [Invitrogen]; 1:200 Alexa 488 anti-chicken [Invitrogen]). Following three additional 10-min washes in PBST, sections were mounted onto micro slides (VWR International, LLC). Vectashield Hart Set Mounting Medium with DAPI (Vector Laboratories, Inc) was applied, slides were cover slipped, and allowed to dry overnight.

### Cell Quantification

Only animals that had accurate bilateral injections were selected for quantification. Fluorescent images were acquired using a confocal microscope (Zeiss LSM800, Germany) at the 20X objective. For cfos quantification, all animals were sacrificed 90 minutes post-behavior for immunohistochemical analyses. The number of EYFP, TdTomato, and *c-Fos*-immunoreactive neurons in the vCA1 were counted to measure the number of active cells in the respective area. 3– 5 coronal slices (spaced 50 um from each other) per mouse of which the means were taken; each N value contains 3-5 image quantification for 3-5 mice. The number of eYFP-positive, TdTomato-positive, and *c-Fos*-positive, and DAPI-positive cells in a set region of interest (ROI) were quantified manually across two different experimenters with ImageJ (https://imagej.nih.gov/ij/). The size of the ROI was standardized across brains, animals, experiments, and groups. The two counters were blind to experimental and control groups. To calculate the percentage of tagged cells we counted the number of fluorescently positive cells and divided them by the total number of DAPI cells. Overlaps were counted as a percentage over DAPI as well, for example, cells that were positive for both EYFP and TdTomato were counted over DAPI. Venn Diagram charts are representative graphs of the proportion of EYFP+, TdTomato+, and cfos+ cells normalized to 100% not to DAPI.

### Fluorescence Intensity Calculations

50 micron coronal sections were stained with chicken anti GFP (1:1000) and rabbit anti RFP (1:1000) as described above. Fluorescent images were taken at 20X magnification using a confocal microscope (Zeiss LSM800, Germany). All laser and imaging settings were maintain consistent across all animals and all brain regions imaged. Regions of interest (ROIs) were maintained consistent across images, sections, and animals. ROIs were identified using landmarks and referencing the Paxinos & Franklin Mouse Brain Atlas. Fluorescence quantification was conducted in ImageJ. After manual selection of the ROI using the selections tool, we set to gather information for area integrated density and mean grey value. Images were analyzed for EYFP and TdTomato separately. Within each image we gathered information about the average fluorescence of the terminals as well as background/baseline levels of a negative region to have a means of normalization. From here we used the following formula for Corrected Total Fluorescence (CTF) = integrated density – (area of selected ROI X mean fluorescence of baseline/background readings).^44–46^ This gave us the arbitrary units for average fluorescence intensity of our EYFP vs TdTomato terminals of interest.

### Image Integrity

Acquired image files (.czi) were opened in ImageJ. Processing of images in Figure 1 involved maximizing intensity, removing outlier noise, and adjusting contrast of images.

### Data Analysis

Data were analyzed using Prism (GraphPad Software, La Jolla, CA) and Statistica 13 data analysis software (TIBCO Software, Inc., Palo Alto, CA). Data were analyzed using paired t-tests (two factors), unpaired t-tests, one-way or two-way ANOVAs with repeated measures ANOVAs (more than two factors), where appropriate. Post-hoc analyses(Tukey’s multiple comparisons test) were used to characterize treatment and interaction effects, when statistically significant alpha set at p< 0.05, two-tailed). Statistical analyses are reported in figure captions.

### Generation of Single Cell Suspension from Mouse Hippocampal Tissue

Five-week old male mice labeled with ChR2-YFP transgene^1^ after conditioning were euthanized by isoflurane. Mouse brains were rapidly extracted, and the hippocampal regions were isolated by microdissection. Eight mice were pooled by each experimental condition. Single cell suspension was prepared according to the guideline of Adult Brain Dissociation Kit (Miltenyi Botec, Cat No: 13-107-677). Briefly, the hippocampal samples were incubated with digestion enzymes in the C Tube placed on the gentleMACS Octo Dissociator with Heaters with gentleMACS Program: 37C_ABDK_01. After termination of the program, the samples were applied through a MACS SmartStrainer (70 μm). Then a debris removal step and a red blood cell removal step were applied to obtain single cell suspension.

### Isolation of YFP-positive single cell by FACS

The single cell suspension was subject to a BD FACSAria cell sorter according to the manufacturer’s protocol to isolate EYFP-single cell population.

### Preparation of RNA-seq library

The RNA of FACS isolated YFP-positive cells was extracted by using Trizol (Life Technologies) followed by Direct-zol kit (Zymo Research) according to manufacturer’s instructions. Then the RNA-seq library was prepared using SMART-Seq® v4 Ultra® Low Input RNA Kit (TaKaRa).

### Analysis of RNA-seq data

The resulting reads from Illumina had good quality by checking with FastQC^63^. The first 40 bp of the reads were mapped to mouse genome (mm10) using STAR^29^, which was indexed with Ensembl GRCm38.91 gene annotation. The read counts were obtained using featureCounts^30^ function from Subread package with unstranded option. Reads were normalized by library size and differential expression analysis based on negative binomial distribution was done with DESeq2^52^. Genes with FDR-adjusted p-value less than 0.00001 plus more than 4-fold difference were considered to be differentially expressed. Raw data along with gene expression levels are being deposited to NCBI Gene Expression Omnibus. For Gene Ontology analysis, we ranked protein-coding genes by their fold changes and used as the input for GseaPreranked tool^53^. Based on the GSEA outputs, the dot plots were created with a custom R script. The pathways with FDR-adjusted p-value less than 0.25 were considered to be enriched. GeneMANIA network were analyzed by GeneMANIA^54^. The max resultant genes were 20 and the max resultant attributes were 10. All networks were assigned an equal weight. The network plots were generated by Cytoscape^55^.

### Analysis of RRBS data

Raw reads were trimmed using cutadapt^56^ and then mapped to mouse genome (mm10) using Bismark^62^ Mcomp module from MOABS^57^ was used to call the differently methylated cytosines (DMCs) and regions (DMRs). Only Cytosines covered with at least five reads were used for further analysis. Differences in DNA methylation levels greater than 0.2 and P value from Fisher Exact Test greater than 0.5 were considered as DMCs. The differently methylated regions at least included two DMCs, and the max distance between two DMCs was 300 bp. Homer^58^ software was used for DMCs and DMRs annotation. GO analysis was done by R package clusterProfiler^59^.

### Sampling strategy

Subjects were randomly assigned to groups. No statistical methods were used to determine sample size; the number of subjects per group were based on those in previously published studies and are reported in figure captions.

### Reproducibility

Behavioral experiments were replicated at least three times with three different experimenters. The first experiments were run at a different institution and the last two replications were run at Boston University. Not only did the behavioral findings stand across experimenters (1 male, 2 females), but it stood across institutions as well. Sequencing data was replicated twice as well to confirm original findings. All behavioral and cell counters and scorers were blinded to experimental and control groups.

### Slice electrophysiology recording

FosCRE-ERT2 male mice were injected with a 1:1 ratio of AAV-CAG-FLEX-TdTomato + AAV-cfos-tTA+AAv-TRE-EYFP into the ventral hippocampus. Animals were counterbalanced for the experiment where half received appetitive tagging with 4OHT and aversive tagging with Dox and the other half received aversive tagging with 4OHT and appetitive tagging with Dox. Coronal slices of the ventral hippocampus were prepared from previously injected animals, 3-5 days post tagging experience. Animals were deeply anesthetized with isoflurane anesthesia, decapitated and the brains were removed. Brain slices were prepared in oxygen perfused sucrose-substitute artificial cerebrospinal fluid (ACSF). Solution contained: 185 mM NaCl, 2.5 mM KCl, 1.25 mM NaH_2_PO_4_, mM MgCl_2_, 25 mM NaHCO_3_, 12.5 mM glucose, and 0.5 mM CaCl_2_. 400 µm slices were cut via Leica VT 1200 (Leica Microsystems) and incubated in 30°C ACSF consisting of 125 mM NaCl, 25 mM NaHCO_3_, 25 mM D-glucose, 2 mM KCl, 2 mM CaCl_2_, 1.25 mM NaH_2_PO_4_, and 1 mM MgCl_2_, for 20 minutes and then cooled to room-temperature. Two-photon imaging system (Thorlabs Inc.) was used to distinguish appetitive and aversive cells. Imaging system is equipped with a mode-locked Ti:Sapphire laser (Chameleon Ultra II; Coherent) which was set to wavelengths between 920 nm to 950 nm in order to excite Alexa Fluor 488 and 568, tdTomato and EYFP fluorophores using a 20x, NA 1.0 objective lens (Olympus). In order to ensure differences between the two populations is not attributable to the type of virus used for tagging the neurons, two groups of animals were prepared. In one group appetitive memories tagged with tdTomato and aversive ones with EYFP and in the other one vice versa. Patch-clamp electrodes with 0.6–1 μm tips were pulled via horizontal puller (Sutter Instruments), and pipette resistance was recorded between 4 and 6 MΩ. Pipettes were filled with intracellular fluid containing: 120 mM K-gluconate, 20 mM KCl, 10 mM HEPES, 7 mM diTrisPhCr, 4 mM Na2ATP, 2 mM MgCl_2_, 0.3 mM Tris-GTP, and 0.2 mM EGTA; buffered to pH 7.3 with KOH. 0.15% weight/volume of Alexa Fluor (Thermo Fisher Scientific) 488 hydrazide (for recording from tdTomato cells) or 567 (for recording from EYFP cells) were added to visualize the recording pipette under the imaging system. Prior to breaking into the cells, pipette capacitance neutralization and bridge balance compensation were performed. Data was acquired using Multiclamp 700B (Molecular Devices) and a Digidata 1550 (Molecular Devices) with sampling rate of 10kHz.

### Electrophysiology data analysis

Brain slice electrophysiology data analysis was performed in python with custom written scripts using pyABF package (http://swharden.com/pyabf/). For spike shape analysis, spike onsets were identified using their first derivative, corresponding voltage was called spike onset. Spike half-width indicates the time between the two halves of peak voltage. For dynamic analysis of spiking properties stepwise current injections were performed. Firing threshold is the voltage at which a neuron fires at least a single spike, and firing onset is the corresponding current. FI gain was calculated as change in the firing frequency from the lowest to the highest frequency of firing divided by current injected. Adaptation ratio was obtained by dividing the mean inter spike intervals (ISIs) during the last 200 ms of spiking by the mean ISIs during the first 200 ms of spiking. Comparison between appetitive and aversive memory cells were done both within a single slice, within an animal as well as across animals to control for potential differences between animals or slices. Since the results in all three situations were similar, the pooled data for comparison of different groups d’Agostino-Pearson K2 test was used to determine normality of data. Based on the normality results, either independent t-test (normal distribution) or Mann-Whitney-Wilcoxon test two-sided with Bonferroni correction were used.

## Acknowledgements

We thank Dr. Joshua Sanes and his lab at the Center for Brain Science, Harvard University, for providing laboratory space within which the initial experiments were conducted, especially the early members of the lab, Emily Merfeld, Emily Doucette, Stephanie Grella, and Joseph Zaki. Further we thank the Center for Brain Science Neuroengineering core for providing technical support, and the Society of Fellows at Harvard University for their support. This work was supported by an NIH Early Independence Award (DP5 OD023106-01), an NIH Transformative R01 Award, a Young Investigator Grant from the Brain and Behavior Research Foundation, a Ludwig Family Foundation grant, the McKnight Foundation Memory and Cognitive Disorders award, and the Center for Systems Neuroscience and Neurophotonics Center at Boston University.

## Author Contributions

M.S. and O.M. conducted and analyzed histology. M.S., O.M., and E.R. ran and analyzed optogenetic experiments. B.Y., X. G. and X.S.L. performed RNA-seq and Reduced Representation Bisulfite sequencing and corresponding analyses. B.R. and F.R.F. conducted in vitro physiological experiments and analyses. M.S., O.M., X.S.L. and S.R. designed the project. M.S. X.S.L. and S.R. wrote the manuscript. All authors edited and commented on the manuscript.

## Extended Data Figures

**Extended Data Fig 1:**
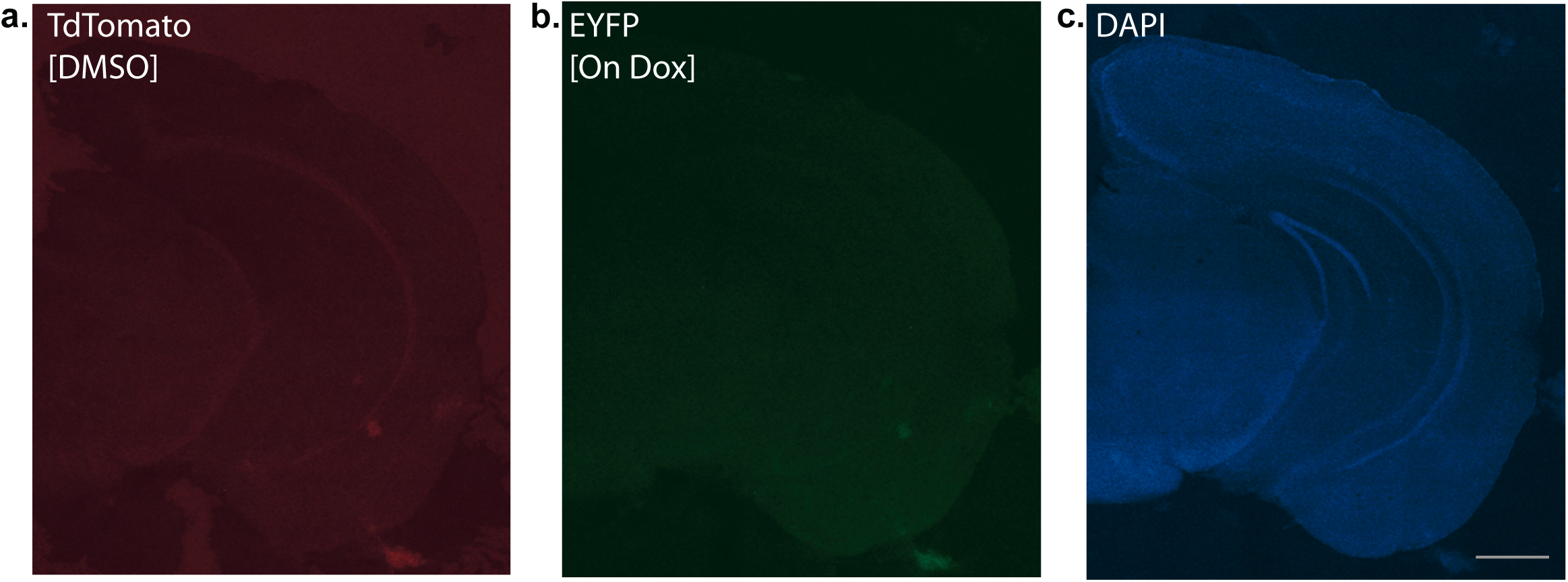
Representative image of TRAP2 ‘leakiness’ in vHPC. **a,** Representative image of TdTomato expression in mouse with DMSO injection instead of 4-OHT. **b,** EYFP expression in the same mouse while remaining on DOX continuously. **c,** DAPI.

**Extended Data Figure 2:**
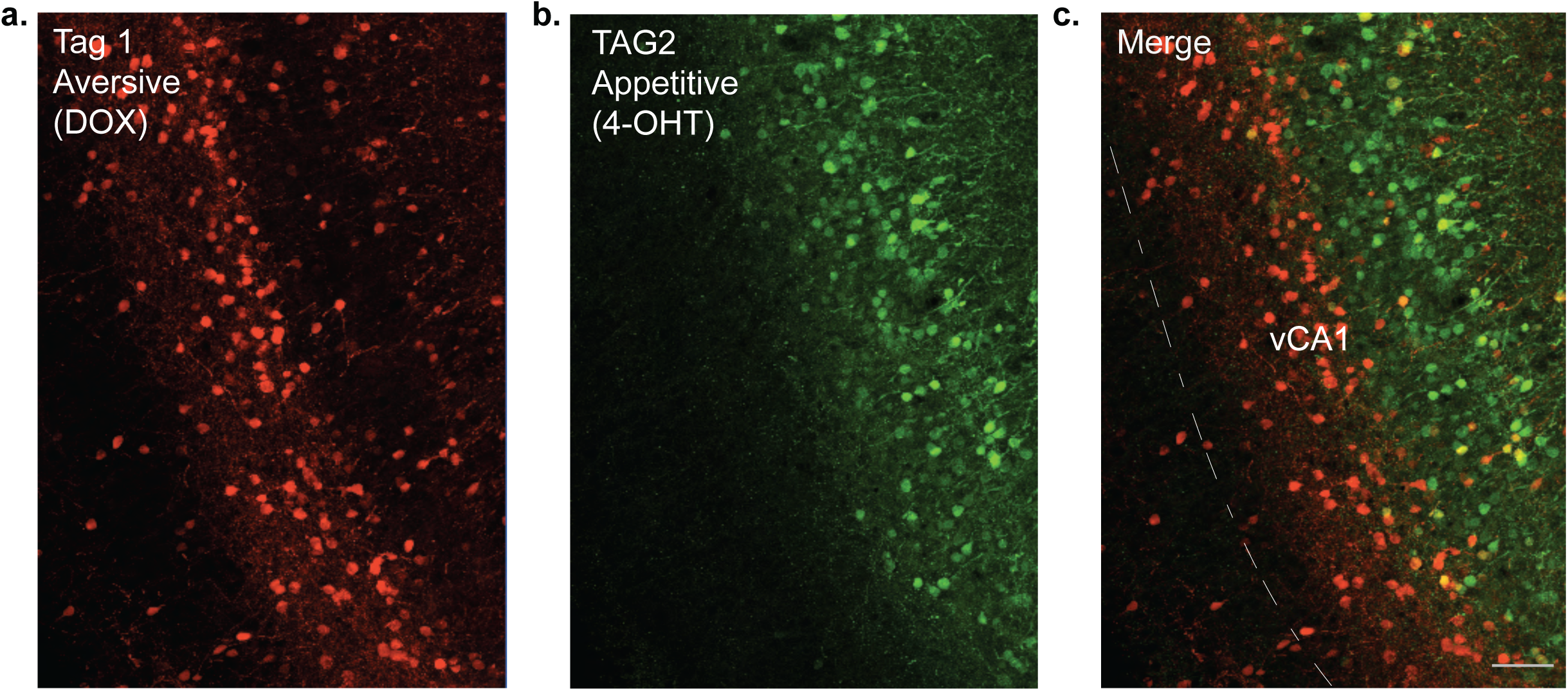
Anterior-posterior vCA1 segregation of appetitive and aversive engrams occurs regardless of experience timing. Here, the mouse was first taken off dox and experienced four foot shocks at 2 seconds each, 4 times. 48 hours later, the mouse was allowed to interact with a female mouse to tag the appetitive experience with 4-OHT. As shown in figures a, b, and c, we still see a clear segregation of appetitive and aversive engrams across vCA1, similar to the segregation shown in Fig 1c-d.

**Extended Data Fig 3:**
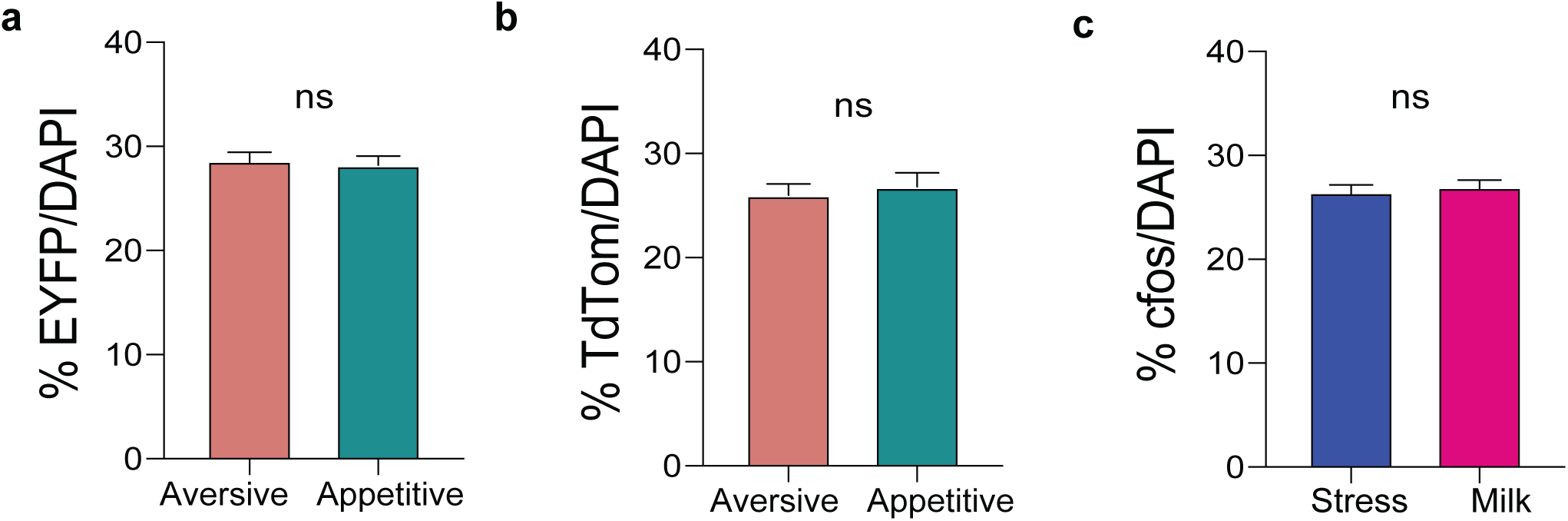
Cell recruitment based on valence type and cfos overlap measures. **a** No significant (ns) difference in % overlap of EYFP over DAPI for shock vs female tags. **b** % No significant (ns) difference in % overlap of TdTomato over DAPI for shock vs female tags. **c** No significant (ns) difference in % overlap of cfos over DAPI for shock vs female tags.

**Extended Data Fig 4:**
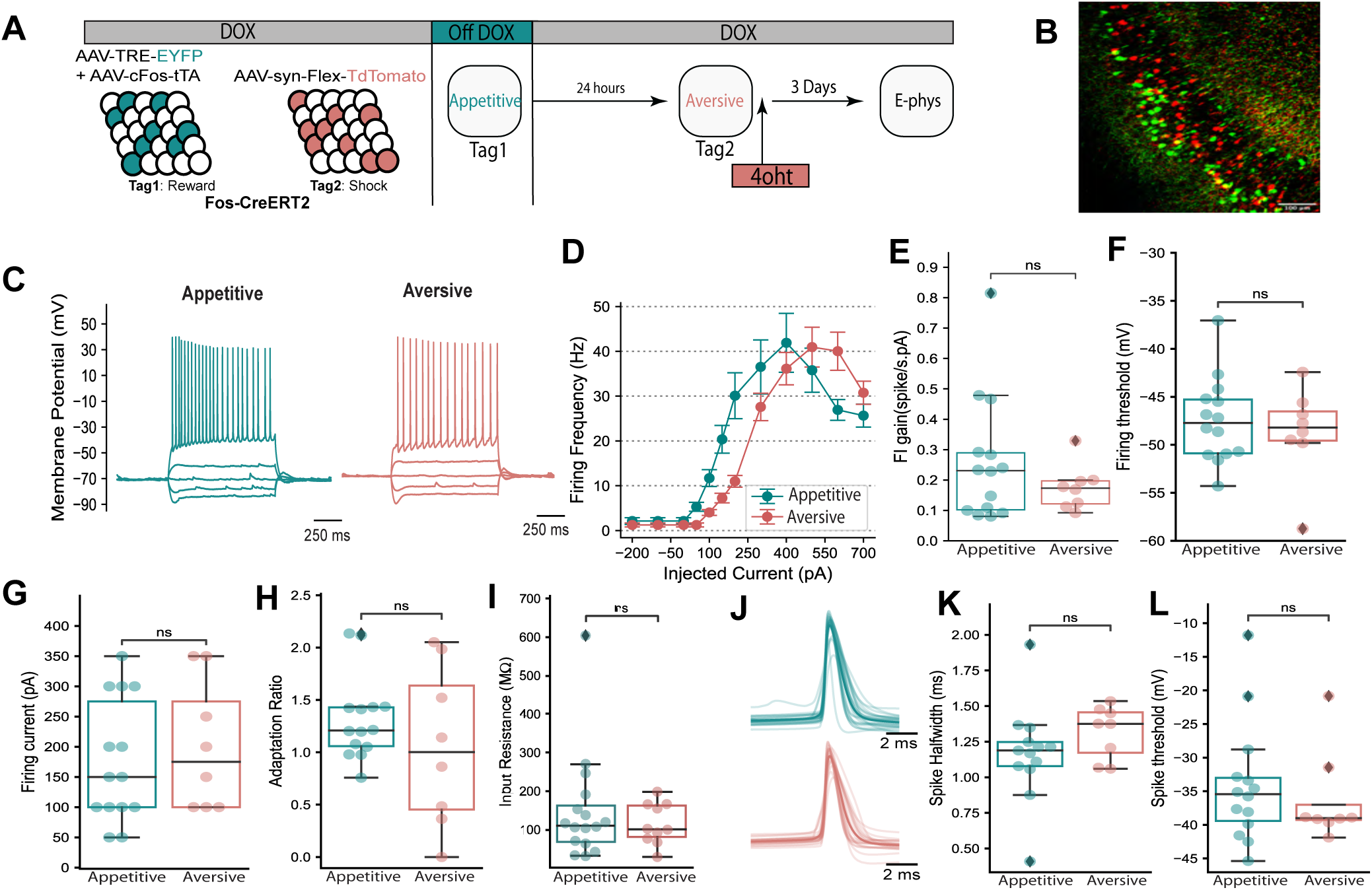
Aversive and appetitive valenced cells in the ventral hippocampus share similar electrophysiological characteristics. **a,** Fos-CreERT2 were injected with a virus cocktail of AAV-Flex-DIO-TdTomato, AAV9-c-Fos-tTA and AAV9-TRE-EYFP to tag either aversive or appetitive experiences within the same animal for slice electrophysiology. **b,** Representative image of cells processing a aversive (red) or appetitive (green) experience. **c,** Representative voltage responses to -100, -50, 0, 50 and 150 pA of step current injection in aversive (red) and appetitive (green) engram cells. **d,** Average firing frequency corresponding to current injection (FI curve) (aversive engrams: n=10, appetitive engrams: n=17). Error bars correspond to standard error and no point of the curve was significantly different between the two populations. **e-g**, Suprathreshold characterization current-voltage responses of tagged hippocampal neurons. E, FI gain is the slope of the linear part of the FI curve shown in D, calculated as the change in firing frequency from the minimum firing to maximum firing divided by the injected current. **h-i**, Voltage and current step at which the neuron had at least one spike. h Adaptation ratio is calculated as the mean inter spike interval (ISI) during the last 200 ms of spiking divided the mean ISI during the first 200 ms of spiking during a current step corresponding to 20 Hz firing rate. i, Input resistance for each neuron, calculated by averaging 25 trials for change in voltage divided by 25 pA injected while holding the cell at -70 mV. **j-l**, Analysis of spike shape of cells tagged by aversive or appetitive experiences. **j**, Spike traces from all the aversive and appetitive cells (Bold trace corresponds to average trace shape). K, Spike half width calculated as the time duration between two midpoints of the spikes. **l**, Deflection point of spike as identified by its first derivative. P-values were calculated based on independent t-test or Mann-Whitney test depending on normality of data.

**Extended Data Fig. 5:**
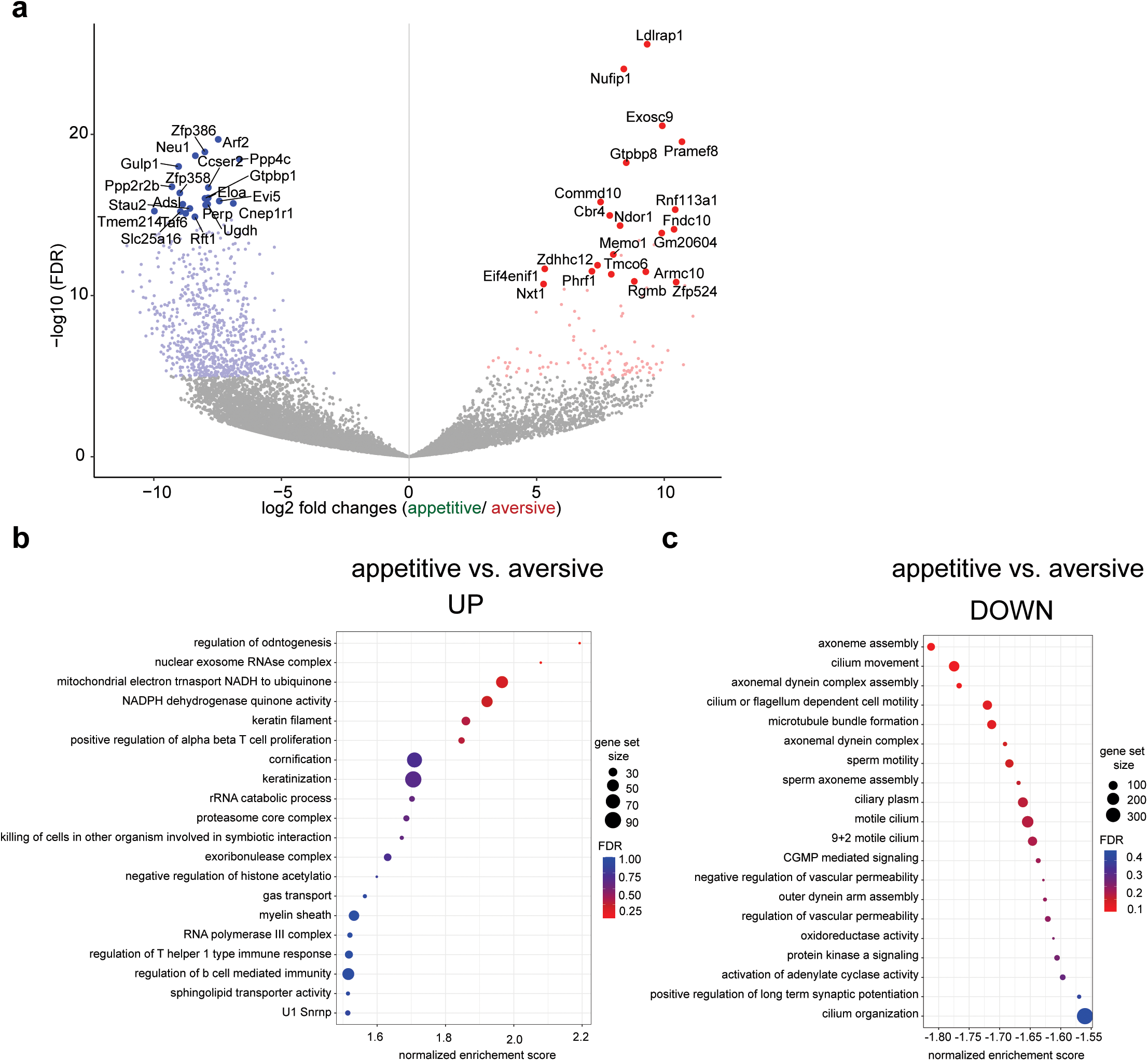
RNA-seq analysis of hippocampus cells processing appetitive and aversive memory engrams. **a,** A Volcano plot with the relative fold change of gene expression in log 2 ratio and the FDR-adjusted p-value in log 2 ratio as X and Y axis showing the up and down regulated genes in appetitive vCA1 cells compared to aversive vCA1 cells. Up or down regulated genes with FDR adjusted p-value less than 1e-5 plus at least four-fold differences were highlighted in either red or blue respectively. The gene names were displayed for the top 20 most significant protein coding genes as sorted by Wald statistics. **b,** Gene Set Enrichment Analysis of the up regulated pathways in appetitive engrams compared to aversive engrams using Gene Ontology (GO) gene sets. The size of the dot represents the gene count and the color of the dot indicates the FDR. Any pathway within a FDR smaller than 0.25 is considered as significantly enriched. **c,** GO analysis of down regulated pathways in appetitive engrams compared to aversive engrams.

**Extended Data Fig. 6:**
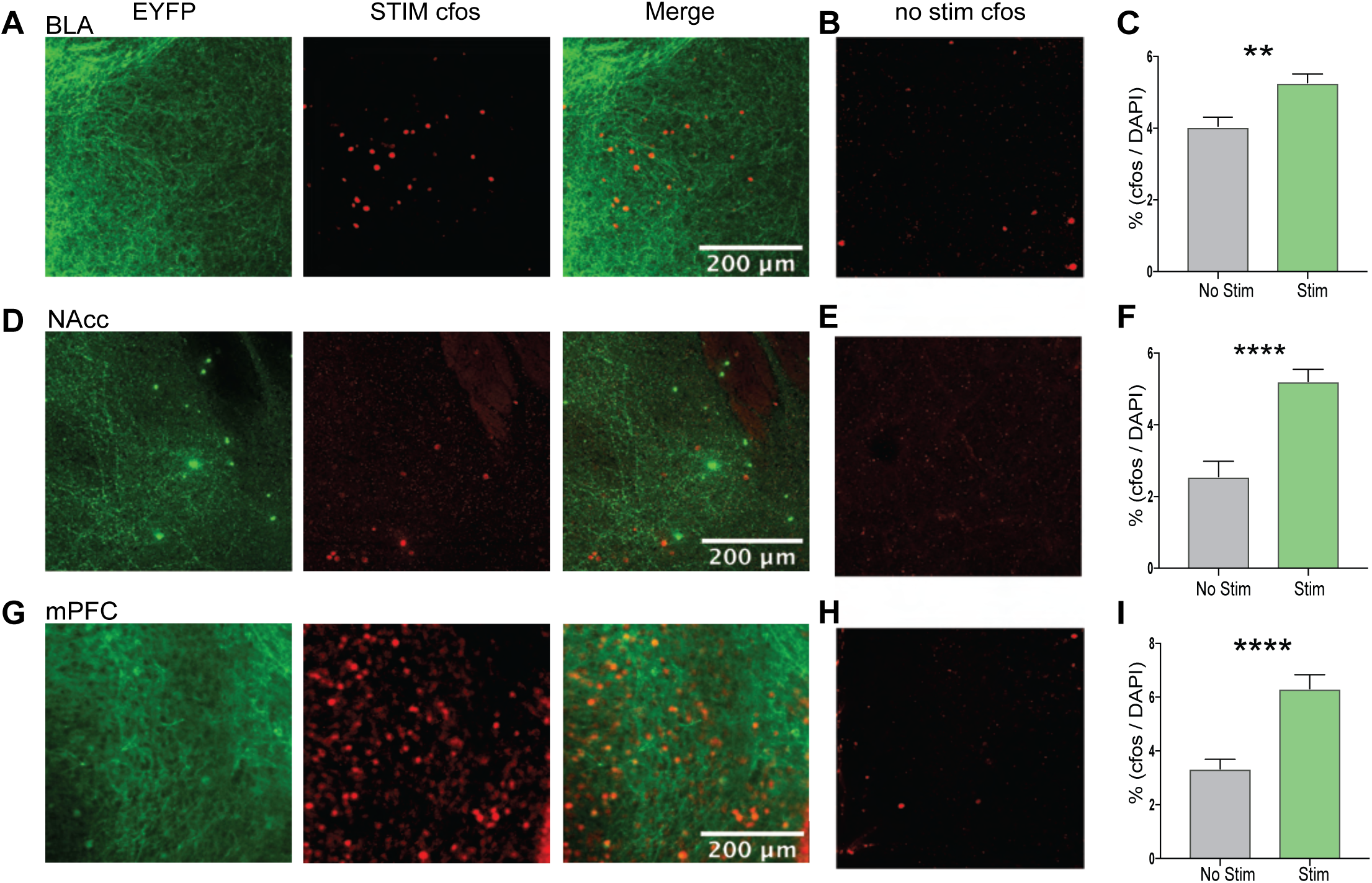
Terminal-specific optogenetic stimulation of vCA1 outputs leads to increased cFos in target areas. **a, d, g,** Representative images of ChR2-EYFP in BLA, NAc, and PFC terminals, respectively, and cFos expression after optical reactivation. **b, e, h,** Representative images of cFos without optogenetic reactivation of corresponding vCA1 terminals. **c, f, i,** Percent cFos/DAPI of BLA, NAc, and PFC non-stim vs stim groups (**P =0.0014, ****P < 0.0001 unpaired student’s t-test)

**Extended Data Fig. 7:**
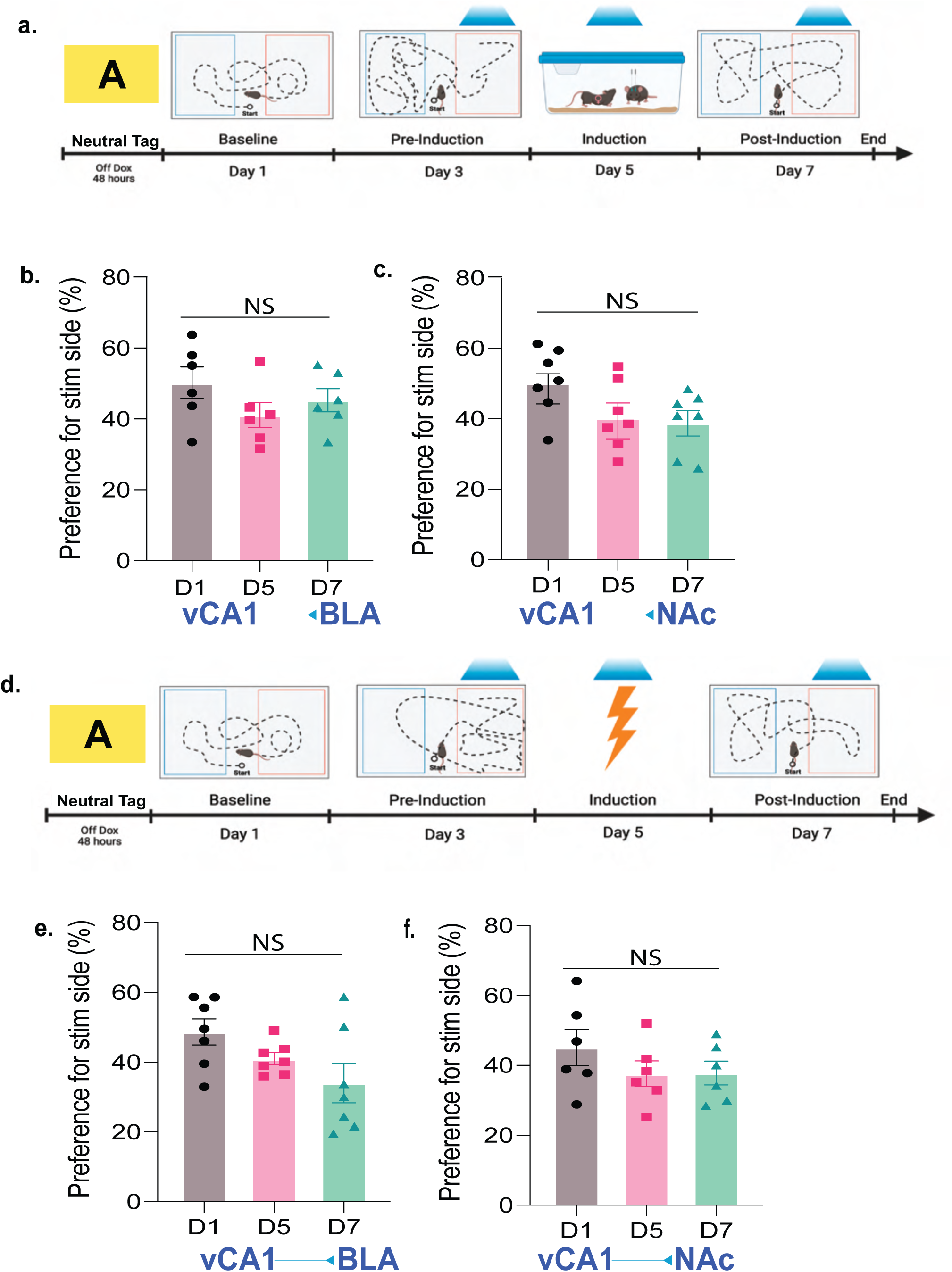
Activating cells processing a neutral memory does not lead to changes in avoidance or preference. **a,** Behavioral schedule in which vCA1 terminals are tagged in a neutral homecage experience (depicted as a yellow square during the off Dox period), followed by the induction protocol in which these outputs are activated concurrently during a appetitive experience in the BLA or NAc. Mice were injected with a virus cocktail consisting of cFos-tTa + TRE-ChR2-EYFP. **b-c,** Percent preference for the stimulation side when optically activating vCA1 terminals over the **b,** BLA or **c,** NAc at 20Hz during baseline (D1), pre-induction (D2), and post-induction (D3) tests. **d,** Behavioral schedule in which vCA1 terminals are tagged in a neutral homecage experience and during induction are reactivated during a aversive (shock) experience. e-f Percent preference for the stimulation side, neutral to aversive, when optically activating vCA1 terminals over the **e,** BLA or **f,** NAc during D1, D5, and D7. NS = not significant; repeated measures one-way ANOVA followed by Tukey’s multiple comparison test).

**Extended Data Fig. 8:**
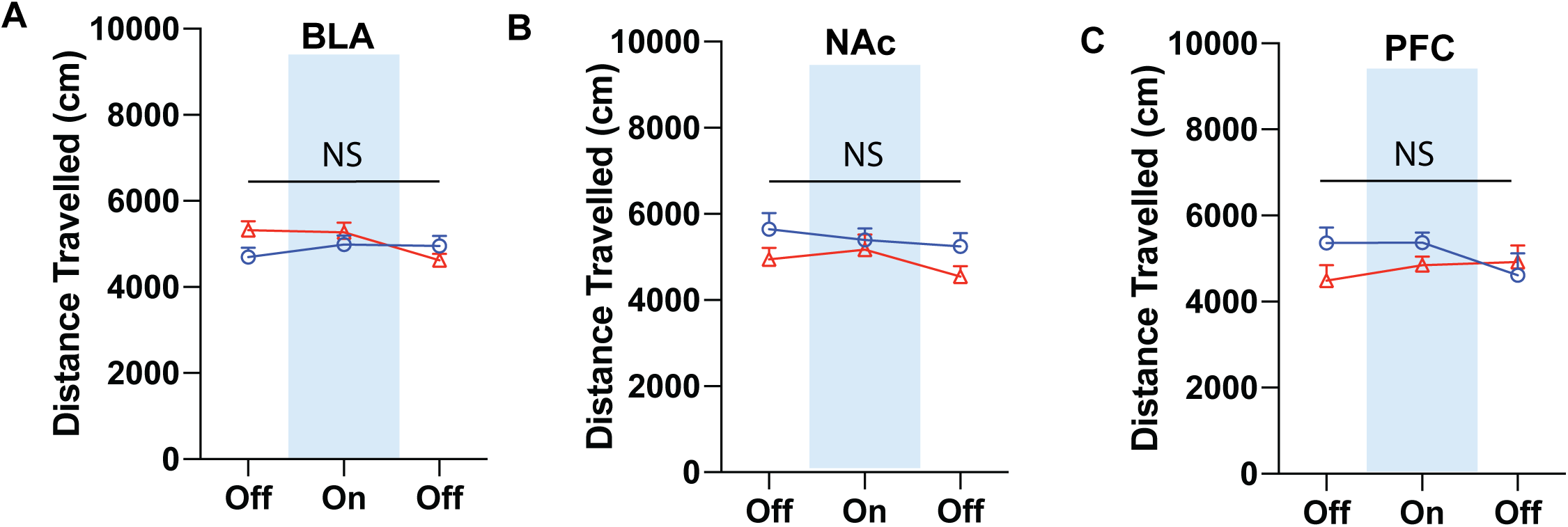
Terminal-specific optogenetic stimulation of vCA1 outputs did not affect distance travelled in all groups. Stimulation of vCA1 terminals over the **a,** BLA (n=8), **b,** NAc (n=10), or **c,** PFC (n=9), had no effect on the distance travelled between light on or off epochs and across pre-induction or post-induction timepoints.

