## Supplementary material for "Hippocampal cells multiplex positive and negative engrams": S1

**Table S1: Top 20 up-regulated and downregulated genes in negative vs. neutral cells**

| SYMBOL | GENE NAME |
|---|---|
| **Upregulated** | |
| Ccl4 | chemokine (C-C motif) ligand 4 |
| Ccl3 | chemokine (C-C motif) ligand 3 |
| Ccl12 | chemokine (C-C motif) ligand 12 |
| Ucp3 | uncoupling protein 3 (mitochondrial, proton carrier) |
| Slitrk2 | SLIT and NTRK-like family, member 2 |
| Etnppl | ethanolamine phosphate phospholyase |
| Ccl7 | chemokine (C-C motif) ligand 7 |
| Adgrb2 | adhesion G protein-coupled receptor B2 |
| Rnf225 | ring finger protein 225 |
| Cd48 | CD48 antigen |
| Fam198a | golgi associated kinase 1A |
| Cst7 | cystatin F (leukocystatin) |
| Drp2 | dystrophin related protein 2 |
| Gli1 | GLI-Kruppel family member GLI1 |
| Tnf | tumor necrosis factor |
| Luzp2 | leucine zipper protein 2 |
| Kcnj9 | potassium inwardly-rectifying channel, subfamily J, member 9 |
| Cxcl2 | chemokine (C-X-C motif) ligand 2 |
| Kcnj16 | potassium inwardly-rectifying channel, subfamily J, member 16 |
| Myoc | myocilin |
| **Downregulated** | |
| Ldlrap1 | low density lipoprotein receptor adaptor protein 1 |
| Rbpms2 | RNA binding protein with multiple splicing 2 |
| Fbxo30 | F-box protein 30 |
| Dvl3 | dishevelled segment polarity protein 3 |
| Gtpbp8 | GTP-binding protein 8 (putative) |
| Usb1 | U6 snRNA biogenesis 1 |
| Mrps27 | mitochondrial ribosomal protein S27 |
| Slc16a11 | solute carrier family 16 (monocarboxylic acid transporters), member 11 |
| Wdr31 | WD repeat domain 31 |
| Mark1 | MAP/microtubule affinity regulating kinase 1 |
| Lrrc41 | leucine rich repeat containing 41 |
| Osbpl5 | oxysterol binding protein-like 5 |
| Matn2 | matrilin 2 |
| Pramef8 | Putative PRAME Family Member 24 |
| Ccdc17 | coiled-coil domain containing 17 |
| Exosc9 | exosome component 9 |
| Gprc5c | G protein-coupled receptor, family C, group 5, member C |
| Ttll1 | tubulin tyrosine ligase-like 1 |
| Gbe1 | glucan (1,4-alpha-), branching enzyme 1 |
| Fance | Fanconi anemia, complementation group E |
