## Supplementary material for "Hippocampal cells multiplex positive and negative engrams": S2

**Table S2: Top 20 upregulated and downregulated genes in positive vs. neutral cells**

| SYMBOL | GENE NAME |
|---|---|
| **Upregulated** | |
| Ccl4 | chemokine (C-C motif) ligand 4 |
| Ccl3 | chemokine (C-C motif) ligand 3 |
| Ccl12 | chemokine (C-C motif) ligand 12 |
| Etnppl | ethanolamine phosphate phospholyase |
| Papss2 | 3'-phosphoadenosine 5'-phosphosulfate synthase 2 |
| Grik2 | glutamate receptor, ionotropic, kainate 2 (beta 2) |
| Csmd2 | CUB and Sushi multiple domains 2 |
| Igdcc4 | immunoglobulin superfamily, DCC subclass, member 4 |
| F3 | coagulation factor III |
| Kcnc4 | potassium voltage gated channel, Shaw-related subfamily, member 4 |
| Gria2 | glutamate receptor, ionotropic, AMPA2 (alpha 2) |
| Rbp4 | retinol binding protein 4, plasma |
| Slc1a2 | solute carrier family 1 (glial high affinity glutamate transporter), member 2 |
| Tnf | tumor necrosis factor |
| Phactr3 | phosphatase and actin regulator 3 |
| Gli1 | GLI-Kruppel family member GLI1 |
| Scg3 | secretogranin III |
| Hrh1 | histamine receptor H1 |
| Ftmt | ferritin mitochondrial |
| Pdzph1 | PDZ and pleckstrin homology domains 1 |
| **Downregulated** | |
| Perp | PERP, TP53 apoptosis effector |
| Hemk1 | HemK methyltransferase family member 1 |
| Ppp4c | protein phosphatase 4, catalytic subunit |
| Hprt | hypoxanthine guanine phosphoribosyl transferase |
| Pcolce | procollagen C-endopeptidase enhancer protein |
| Sdf2l1 | stromal cell-derived factor 2-like 1 |
| Arf2 | ADP-ribosylation factor 2 |
| Trpv4 | transient receptor potential cation channel, subfamily V, member 4 |
| Evi5 | ecotropic viral integration site 5 |
| Fbxw11 | F-box and WD-40 domain protein 11 |
| Spint2 | serine protease inhibitor, Kunitz type 2 |
| Slc31a1 | solute carrier family 31, member 1 |
| Rfng | RFNG O-fucosylpeptide 3-beta-N-acetylglucosaminyltransferase |
| Tmem214 | transmembrane protein 214 |
| Eloa | elongin A |
| Ttc9 | tetratricopeptide repeat domain 9 |
| Ugdh | UDP-glucose dehydrogenase |
| Acd | adrenocortical dysplasia |
| Adsl | adenylosuccinate lyase |
| Zdhhc1 | zinc finger, DHHC domain containing 1 |
