## Supplementary material for "Hippocampal cells multiplex positive and negative engrams": S3

**Table S3: Top 20 upregulated and downregulated genes in positive vs. negative cells**

| SYMBOL | GENE NAME |
|---|---|
| **Upregulated** | |
| Ldlrap1 | low density lipoprotein receptor adaptor protein 1 |
| Nufip1 | nuclear fragile X mental retardation protein interacting protein 1 |
| Exosc9 | exosome component 9 |
| Pramef8 | Putative PRAME Family Member 24 |
| Gtpbp8 | GTP-binding protein 8 (putative) |
| Commd10 | COMM domain containing 10 |
| Rnf113a1 | ring finger protein 113A1 |
| Cbr4 | carbonyl reductase 4 |
| Ndor1 | NADPH dependent diflavin oxidoreductase 1 |
| Fndc10 | fibronectin type III domain containing 10 |
| Gm20604 | predicted gene 20604 |
| Memo1 | mediator of cell motility 1 |
| Zdhhc12 | zinc finger, DHHC domain containing 12 |
| Eif4enif1 | eukaryotic translation initiation factor 4E nuclear import factor 1 |
| Phrf1 | PHD and ring finger domains 1 |
| Armc10 | armadillo repeat containing 10 |
| Tmco6 | transmembrane and coiled-coil domains 6 |
| Rgmb | repulsive guidance molecule family member B |
| Zfp524 | zinc finger protein 524 |
| Nxt1 | NTF2-related export protein 1 |
| Downregulated | |
| Arf2 | ADP-ribosylation factor 2 |
| Zfp386 | zinc finger protein 386 (Kruppel-like) |
| Neu1 | neuraminidase 1 |
| Ppp4c | protein phosphatase 4, catalytic subunit |
| Gulp1 | GULP, engulfment adaptor PTB domain containing 1 |
| Ppp2r2b | protein phosphatase 2, regulatory subunit B, beta |
| Ccser2 | coiled-coil serine rich 2 |
| Zfp358 | zinc finger protein 358 |
| Eloa | elongin A |
| Gtpbp1 | GTP binding protein 1 |
| Evi5 | ecotropic viral integration site 5 |
| Cnep1r1 | CTD nuclear envelope phosphatase 1 regulatory subunit 1 |
| Ugdh | UDP-glucose dehydrogenase |
| Adsl | adenylosuccinate lyase |
| Perp | PERP, TP53 apoptosis effector |
| Stau2 | staufen double-stranded RNA binding protein 2 |
| Tmem214 | transmembrane protein 214 |
| Slc25a16 | solute carrier family 25 (mitochondrial carrier, Graves disease autoantigen), member 16 |
| Taf6 | TATA-box binding protein associated factor 6 |

| | |
|---|---|
| Rft1 | RFT1 homolog |
