## Supplementary material for "Hippocampal cells multiplex positive and negative engrams": S4

**Table S4 GeneMANIA Network**

**Shared protein domains**

A-amylase/branching_C
Aminotrans_3
Chemokine_CC_CS
Chemokine_IL8-like_dom
Cyclic_Pdiesterase
Dishevelled_fam
Dishevelled_protein_dom
DIX
ExoRNase_PH_dom2
F-box_dom
Fringe
Fumarate_lyase_fam
Glu/Gly-bd
GPCR_2_brain_angio_inhib
Ig_E-set
Interleukin_8-like_sf
Iono_rcpt_met
Iontro_rcpt
IRK_C
K_chnl_inward-rec_Kir
K_chnl_volt-dep_Kv3
KA1_dom
Kir_TM
L-Aspartase-like
Matrilin_cc_sf
Matrilin_coiled-coil_trimer
Mit_uncoupling_UCP-like
Na-dicarboxylate_symporter
Na:dicarbo_symporter_sf
PRibTrfase_dom
RPEL_repeat
Small_mtfrase_dom
TNF
TRPV1-4_channel

**_shared protein domains**

**Links**

Alpha-amylase/branching enzyme, C-terminal all beta
Aminotransferase class-III
CC chemokine, conserved site
Chemokine interleukin-8-like domain
Cyclic phosphodiesterase
Dishevelled family
Dishevelled protein domain
DIX domain
Exoribonuclease, phosphorolytic domain 2
F-box domain
Fringe
Fumarate lyase family
Ionotropic glutamate receptor, L-glutamate and glycine-binding domain
GPCR, family 2, brain-specific angiogenesis inhibitor
Immunoglobulin E-set
Chemokine interleukin-8-like superfamily
Ionotropic glutamate receptor, metazoa
Ionotropic glutamate receptor
Inward rectifier potassium channel, C-terminal
Potassium channel, inwardly rectifying, Kir
Potassium channel, voltage dependent, Kv3
Kinase associated domain 1 (KA1)
Potassium channel, inwardly rectifying, transmembrane domain
L-Aspartase-like
Matrilin, coiled-coil domain superfamily
Matrilin, coiled-coil trimerisation domain
Mitochondrial carrier UCP-like
Sodium:dicarboxylate symporter
Sodium:dicarboxylate symporter superfamily
Phosphoribosyltransferase domain
RPEL repeat
Methyltransferase small domain
Tumour necrosis factor
Transient receptor potential cation channel subfamily V member 1-4
