## Supplementary material for "Hippocampal cells multiplex positive and negative engrams": S5

**Tabel S5. Genome position annotiation of DMCs**

**Genome position of DMCs in Negative vs Neutral**

| Genome position | Number of DMCs | Distribution |
|---|---|---|
| 3' UTR | 27 | 1.39% |
| 5' UTR | 1 | 0.05% |
| Exon | 29 | 1.50% |
| Intergenic | 1008 | 51.99% |
| Intron | 779 | 40.18% |
| Non-coding | 30 | 1.55% |
| Promoter | 33 | 1.70% |
| TTS | 32 | 1.65% |

**Genome position of DMCs in Positive vs Neutral**

| Genome position | Number of DMCs | Distribution |
|---|---|---|
| 3' UTR | 29 | 0.93% |
| 5' UTR | 0 | 0.00% |
| Exon | 89 | 2.86% |
| Intergenic | 1514 | 48.57% |
| Intron | 1305 | 41.87% |
| Non-coding | 68 | 2.18% |
| Promoter | 60 | 1.92% |
| TTS | 52 | 1.67% |

Promoter defined from -1kb to transcription start site(TSS).
TSS: Transcription termination site, defined from -100bp to +1kb
