## Supplementary material for "Hippocampal cells multiplex positive and negative engrams": S6

**Table S6. DMGs**

**DMGs in Negative vCA1 cells (compared with Neutral)**

Mir1970c
Rn45s
Mir7686
Olfr781
Galk1
1700001K19Rik
Hmmr
Pdcd6ip
Gm20752
A730082K24Rik
Ssxb10
Platr9
Sh3pxd2b
Angpt2
Mindy2
Zfp609
Thsd7b
Phlpp1
Ccdc33
Bmper
Nisch
Greb1
Gmppb
Gabrg3
Dcc
Col19a1
Aif1
Mecom
Anks1b
Pea15a
Nyap2
4930467E23Rik
Gm5136
Ykt6
7420701I03Rik
Gm12596
Fam78b
Sox5
Runx1
Cntn5
Cntnap2
March11
Slc16a7

Gm14015
Slitrk2
Mir1952
Lrrc52
Mir6354
Htr1d
a
Lclat1
Gm14725
Map3k7
Hmga2
Ak2
Mir6341
Hey1
Stxbp6
Slc27a6
Gm4632
Gpr165
Shtn1
Nfkbiz
Tmem74
Pou4f3
Sec61b
Tnfsf15
1700022A22Rik
Gm36669
Ripk4
Gm813
Cdh9
Mex3c
Ifnb1
Bambi
Smim10l1
Otud4
Ccdc85a
Slc35b3
Trib2
Asb1
9230019H11Rik
Prr5
Rgs18
Mir7j
Poteg
Tmem139
Zfp275

Cdh6
Hdgfl1
Ccdc85c
Cps1
1700054K19Rik
Ppp1r2-ps3
Gm8096
Sez6
Bloc1s3
Mir7021

**DMGs in Positive vCA1 cells (compared with Neutral)**

Mir1970c
Rn45s
Olfr781
Plcxd1
1700001K19Rik
Zfp955a
Mir1668
Fzd1
1700057G04Rik
Luzp2
Cerk
Gng5
Fos
Nrsn1
Runx2
Cmtr1
Tnk2os
Smad4
Mir879
Gm20319
Trap1
Gm28590
Zfp296
4933402J07Rik
Aopep
8430422H06Rik
Gm20752
Elmo2
Xxylt1
Akap6
4930474N09Rik
Morf4l1-ps1
Opcml
Rtn4rl1
Tmc3
Sh3pxd2b
Tafa4
Dpy19l1
4930505G20Rik
Anxa3
Syt16
Slc38a7
Csgalnact1

Gm6402
Polq
Crocc
Fam129b
St6gal1
Zfp521
Zbtb7c
Mir6896
Acly
Mir378a
Otud7a
Mir3106
Lrrn3
Psapl1
Ift43
Gm5089
Ccdc33
Snx29
Cmip
Bmper
Ppp2r2b
Syde1
Wars2
Psmd1
Gm13205
4930578I06Rik
Dtnbp1
Gtf2a1l
Nisch
Tmod1
Spata5
Evc
Dnah10
Cd244a
Clca1
Suv39h1
Vdac3
Klk1b21
Scn10a
Prim2
Cebpg
4930430F21Rik
4933417O13Rik
Mir592
Gch1

Acte1
Dcc
Col19a1
Slc7a1
Angpt2
Rnf125
Zfp644
Sfi1
Lingo1
Mecom
Pibf1
Sipa1l2
Mir3085
Chchd3
Foxp1
BC031361
Trim68
Itga10
Palld
Cryab
Upp2
Gng2
Vasp
Ica1
Fbxw7
Fgfbp1
Pex5l
Fam78b
Mcf2l
Phtf1os
Wdfy4
Shank2
Runx1
Dbx1
Slc2a9
Popdc2
Pcdh9
Myo5c
Hpse2
Kcnh4
Gtf2i
March4
Acsl6
Cadm1
Crebbp

Bcat1
Cntnap2
Sirpb1a
Lrrc3b
Lclat1
Ugt8a
Stam2
Suclg2
Mlph
Gm12596
Rgs18
4930429B21Rik
Tcf3
Rasl11a
Mterf4
Ppp4r2
Stxbp6
Eno1
Rnf144b
Gm833
Rrp15
1700063A18Rik
Mir6899
Rreb1
Amz1
Trib2
Zfp804a
Diaph2
Slc17a3
Btg4
Nuak1
Mospd2
H60c
Pde1a
1190028D05Rik
Rab3d
BB123696
Olfr381
Gm29683
Gpr26
Atp1a1
Abhd17c
Gm14725
Spaca1
Tmem158

D630023F18Rik
4930563J15Rik
Cdh9
Unc5d
Ftmt
Msl3l2
Zfp770
Shtn1
Mid1ip1
Asprv1
Cacng8
Styk1
Adgrl3
Ccdc85c
Mir3072
Gpcpd1
Actr3
Frem1
Slc35b3
1700028P15Rik
1700017L05Rik
Cenpa
Hmga2
Cps1
Olfm4
Tmsb10
Tmem200c
1700008P02Rik
Art2b
Gm7102
Ppargc1a
Snx10
Dio3os
Lyplal1
Dcdc2c
Scd4
4930470O06Rik
Rasd2
A930031H19Rik
Gm8096
1700049L16Rik
Esp8
Lrrn2
Adam23
Ccng1

Mir6341
4930556N09Rik
Tbl1xr1
Olfm1
Jun
Ankrd16
5930438M14Rik
4933406K04Rik
Ctsm
Basp1
Kcnj2
Calm3
Smarca2
Meis1
Arl6ip6
Poteg
Slc22a16
Arhgap20
Dazl
AA545190
Grk3
Mir6354
Ano5
Nmt2
Gm20740
Ly6d
Bcar3
Foxo3
Ccdc50
E330011O21Rik
9530068E07Rik
Adgrg6
Srbd1
Col15a1
Pin1
6430706D22Rik
Eif2b5
Wrnip1
Tril
Ehmt2
Stox2
Dux
Tmem174

**32 common elements in "Negative DMGs" and "Positive DMGs":**

Dcc
Mir6354
Fam78b
Rn45s
Lclat1
Gm14725
Hmga2
Mir6341
Stxbp6
Shtn1
Mir1970c
Nisch
Gm12596
Gm20752
1700001K19Rik
Cdh9
Col19a1
Angpt2
Runx1
Mecom
Ccdc33
Slc35b3
Olfr781
Trib2
Rgs18
Poteg
Ccdc85c
Cntnap2
Cps1
Sh3pxd2b
Gm8096
Bmper
